# GATA3 and MDM2 are synthetic lethal in estrogen receptor-positive breast cancers

**DOI:** 10.1101/2020.05.18.101998

**Authors:** Gaia Bianco, Mairene Coto-Llerena, John Gallon, Stephanie Taha-Mehlitz, Venkatesh Kancherla, Martina Konantz, Sumana Srivatsa, Hesam Montazeri, Federica Panebianco, Marta De Menna, Viola Paradiso, Caner Ercan, Ahmed Dahmani, Elodie Montaudon, Niko Beerenwinkel, Marianna Kruithof-de Julio, Luigi M. Terracciano, Claudia Lengerke, François-Clément Bidard, Rinath M. Jeselsohn, Elisabetta Marangoni, Charlotte K. Y. Ng, Salvatore Piscuoglio

**Affiliations:** Visceral Surgery and Precision Medicine Research Laboratory, Department of Biomedicine, University of Basel, Basel, Switzerland; Institute of Medical Genetics and Pathology, University Hospital Basel, Basel, Switzerland; Department of Biomedicine, University of Basel and University Hospital Basel, Basel, Switzerland; Department of Biosystems Science and Engineering, ETH Zurich, Basel, Switzerland; Department of Bioinformatics, Institute of Biochemistry and Biophysics, University of Tehran, Tehran, Iran; Department of Biomedical Research, Urology Group, University of Bern, Bern, Switzerland; Laboratory of Preclinical Investigation, Department of Translational Research, Institut Curie Research Center, Paris, France; Department of Medical Oncology, Institut Curie, Saint Cloud, France; Division of Women’s Cancers, Dana-Farber Cancer Institute, Harvard Medical School. Boston. USA; Department for BioMedical Research (DBMR), University of Bern, Bern, Switzerland

**Keywords:** *MDM2*, *GATA3*, breast cancer, nutlins, synthetic lethality, patient-derived organoids, PI3K/Akt/mTOR signaling pathway

## Abstract

Synthetic lethal interactions, where the simultaneous but not individual inactivation of two genes is lethal to the cell, have been successfully exploited to treat cancer. *GATA3* is frequently mutated in estrogen receptor (ER)-positive breast cancers and its deficiency defines a subset of patients with poor response to hormonal therapy and poor prognosis. However, GATA3 is not targetable. Here we show that *GATA3* and *MDM2* are synthetically lethal in ER-positive breast cancer. Depletion and pharmacological inhibition of MDM2 induce apoptosis in *GATA3*-deficient models *in vitro, in vivo* and in patient-derived organoids (PDOs) harboring *GATA3* somatic mutation. The synthetic lethality requires p53 and acts via the PI3K/Akt/mTOR pathway. Our results present MDM2 as a novel therapeutic target in the substantial cohort of ER-positive, *GATA3*-mutant breast cancer patients. With MDM2 inhibitors widely available, our findings can be rapidly translated into clinical trials to evaluate in-patient efficacy.

## Introduction

*GATA3* is mutated in 12-18% of primary and metastatic estrogen receptor (ER)-positive breast cancers, with predominantly frameshift mutations and mutations affecting splice sites (Bertucci et al., 2019; Hoadley et al., 2018). It is the most highly expressed transcription factor in the mammary epithelium (Kouros-Mehr et al., 2006) and has key functions in mammary epithelial cell differentiation (Kouros-Mehr et al., 2006). In breast cancer, GATA3 suppresses epithelial-to-mesenchymal transition (Chou et al., 2013) and acts as a pioneer transcription factor by recruiting other cofactors such as ERα and FOXA1 (Theodorou et al., 2013; Zaret and Carroll, 2011). Its expression level is strongly associated with ERα expression and is diagnostic of the luminal A and luminal B subtypes. Indeed, GATA3 loss has also been strongly linked to poor response to hormonal therapy and poor prognosis (Gulbahce et al., 2013; Liu et al., 2016; Mehra et al., 2005). Therefore, targeting GATA3 deficiency may provide a specific and tailored treatment for a subclass of patients associated with a worse prognosis and relapse.

Synthetic lethality refers to the interaction between genetic events in two genes whereby the inactivation of either gene results in a viable phenotype, while their combined inactivation is lethal (Ashworth and Lord, 2018). It has helped extend precision oncology to targeting genes with loss-of-function mutations by disrupting the genetic interactors of the mutated gene. One such example is the use of poly(ADP-ribose) polymerase (PARP) inhibition in cancers with deficiencies in homologous recombination (Bryant et al., 2005). Recent developments in large-scale perturbation screens have enabled the comprehensive screening of genetic interactions (McDonald et al., 2017) and the systematic analysis of these screens has led to the discovery of further synthetic lethal targets in cancer (Chan et al., 2019; Lieb et al., 2019). In this study, using our recently developed SLIdR (Synthetic Lethal Identification in R) algorithm (Srivatsa et al.), we systematically interrogate the project DRIVE RNAi screen (McDonald et al., 2017) and identify MDM2 as a synthetic lethal interactor of GATA3 in ER-positive breast cancer. We show that inhibition of MDM2 is synthetically lethal in *GATA3*-mutant and GATA3-depleted breast cancer cells. Our findings establish a new approach for targeting GATA3 deficiency in ER-positive breast cancer by pharmacological inhibition of MDM2 using selective small molecules which are currently being evaluated in clinical trials (Jiang and Zawacka-Pankau, 2020).

## Results

### *GATA3* and *MDM2* are synthetic lethal in ER-positive breast cancer

To identify synthetically lethal vulnerabilities of *GATA3* in breast cancer, we analyzed the breast cancer cell line (n=22) data from the large-scale, deep RNAi screen project DRIVE (McDonald et al., 2017) using our recently developed SLIdR algorithm (Srivatsa et al.). SLIdR uses rank-based statistical tests to compare the viability scores for each gene knockdown between the *GATA3*-mutant and *GATA3*-wild type cell lines (**Figure 1A**) and identified *MDM2* as the top gene whose knock-down significantly reduced cell viability in the two *GATA3*-mutant breast cancer cell lines (**Figure 1B-C**). *MDM2* encodes an E3 ubiquitin ligase that inhibits the tumor suppressor p53-mediated transcriptional activation (Momand et al., 1992) and is frequently amplified and overexpressed in human cancers, including breast (Wade et al., 2013).

**Figure 1:**
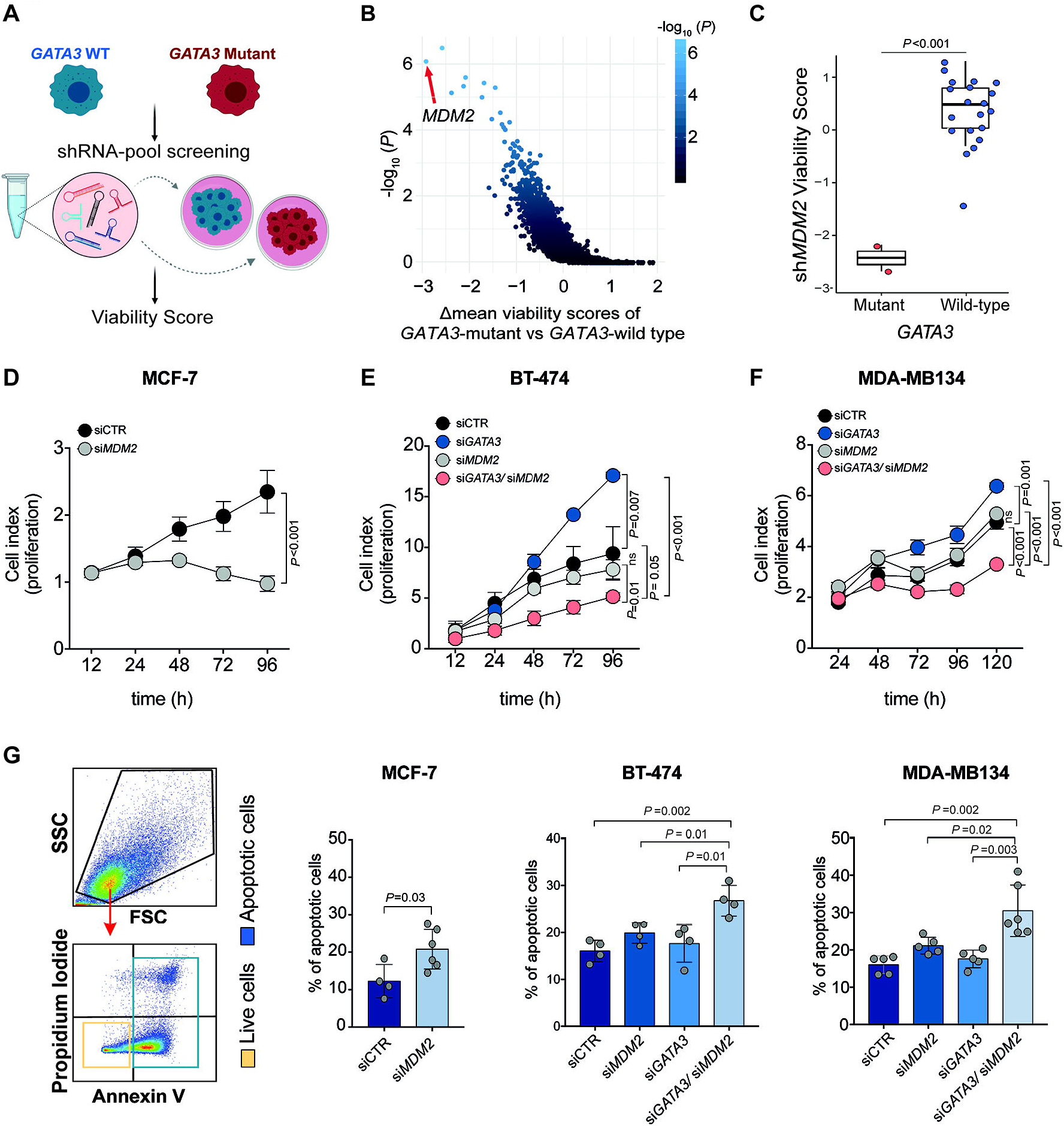
*GATA3* and *MDM2* are synthetic lethal in ER-positive breast cancer. (**A)** Schematic representation of the project DRIVE shRNA screen data used to identify synthetic lethal interactors of *GATA3*. (**B)** SLIdR-derived statistical significance (-log_10_(*P*)) plotted against the difference in the mean viability scores between *GATA3*-mutant and *GATA3*-wild type breast cancer cell lines for each gene knocked-down in the shRNA screen. Middle lines of the boxplots indicate medians. Box limits are first and third quartiles. The whiskers extend to the range. (**C)** Viability scores of *MDM2* knock-down in *GATA3*-mutant and *GATA3*-wild type cell lines. (**D-F)** Proliferation kinetics of (**D)** *GATA3*-mutant MCF-7 transfected with siRNA targeting *MDM2* or control (see also Figure S1A-C), (**E)** *GATA3*-wild type BT-474, (**F)** *GATA3*-wild type MDA-MB134 transfected with siRNA targeting *GATA3, MDM2, GATA3/ MDM2*, or control (see also Figure S1E-F). (**G)** Apoptosis assay using Annexin V and propidium iodide co-staining. From left: gating strategy to define apoptotic (yellow) and live (blue) cells; percentage of apoptotic and live cells upon *MDM2* silencing in MCF-7 (see also Figure S1D); upon silencing of *GATA3* or *MDM2* alone or in combination in BT-474 and MDA-MB134. Data are mean ± s.d. (**D,E,F,G)** n≥4 replicates. Statistical significance was determined for (**D,E,F,G)** by the two-tailed unpaired Student’s t-test.

We first sought to validate the predicted synthetic lethality between *GATA3* and *MDM2* in the ER-positive breast cancer cell line MCF-7, one of the two *GATA3*-mutant cell lines used in the RNAi screen (McDonald et al., 2017). MCF-7 harbors the *GATA3* frameshift mutation p.D335Gfs (Barretina et al., 2012), a loss-of-function mutation that has been recurrently observed in breast cancer patients (Hoadley et al., 2018; Pereira et al., 2016) and leads to a truncated GATA3 protein. Using a siRNA approach, we confirmed that silencing *MDM2* significantly reduced cell proliferation in MCF-7 cells (**Figure 1D, Figure S1A**). *MDM2* siRNA titration analysis showed that the vulnerability induced by *MDM2* inhibition in MCF-7 was dose-dependent and that 50% reduction in *MDM2* expression is sufficient to inhibit proliferation in the presence of *GATA3* mutation (**Figure S1B-C**).

To confirm that the effect of *MDM2* silencing is unequivocally related to GATA3 loss of function and to exclude any gain-of-function effects of the *GATA3* mutation, we assessed the changes in cell proliferation upon single- and dual-silencing of *GATA3* and *MDM2* using siRNA in two ER-positive *GATA3*-wild type breast cancer cell lines; the luminal A (ER+/HER2-) MDA-MB134 and the luminal B (ER+/HER2+) BT-474 (**Figure S1D-E**). Consistent with the oncosuppressor role of *GATA3* in breast cancer (Dydensborg et al., 2009; Yan et al., 2010), *GATA3*-silencing led to a significant increase in cell proliferation in both BT-474 and MDA-MB134 (**Figure 1D-E**). By contrast, dual-silencing of *GATA3* and *MDM2* significantly reduced cell proliferation compared to cells transfected with control siRNA, *GATA3* siRNA or *MDM2* siRNA alone (**Figure 1D-E**).

To determine if *MDM2* silencing was merely inhibiting cell growth or was actively inducing cell death, we assessed apoptosis using Annexin V and propidium iodide co-staining followed by flow cytometry analysis. We observed that *MDM2* silencing significantly induced apoptosis in MCF-7 cells in a dose-dependent manner (**Figure 1G, Figure S1F**). Similarly, *dual-GATA3/MDM2* silencing in BT-474 and MDA-MB134 cells led to 15-20% higher proportion of apoptotic cells than the silencing of the two genes individually (**Figure 1G**), indicating that dual inhibition induced increased apoptosis.

Our results provide evidence that MDM2 is a selected vulnerability in breast cancer with *GATA3*-mutation and/or loss of *GATA3*.

### GATA3 status determines response to MDM2 inhibitor *in vitro*

The selected vulnerability of MDM2 in *GATA3*-deficient ER-positive breast cancers presents MDM2 as an attractive therapeutic target in this patient cohort. To test whether the apoptotic effects of MDM2 inhibition could be achieved using an MDM2 antagonist, we treated the breast cancer cell lines with idasanutlin (RG7388, **Figure S2A**) (Ding et al., 2013; Reis et al., 2016). In the *GATA3*-mutant MCF7 cells, idasanutlin induced cell growth arrest and apoptosis in a dose-dependent manner (**Figure 2A-B**). To assess if idasanutlin was inducing the canonical apoptotic cascade, we assessed the expression of p53, Bax and Bcl-2, together with the canonical markers of apoptosis PARP and cleaved PARP, by immunoblot at 6, 12 and 24 hours post-treatment. Idasanutlin induced an early up-regulation of p53 and MDM2 proteins (Pan et al., 2017), together with the up- and down-regulation of pro- and anti-apoptotic proteins, respectively (**Figure 2C**), leading to the activation of the apoptotic cascade.

**Figure 2:**
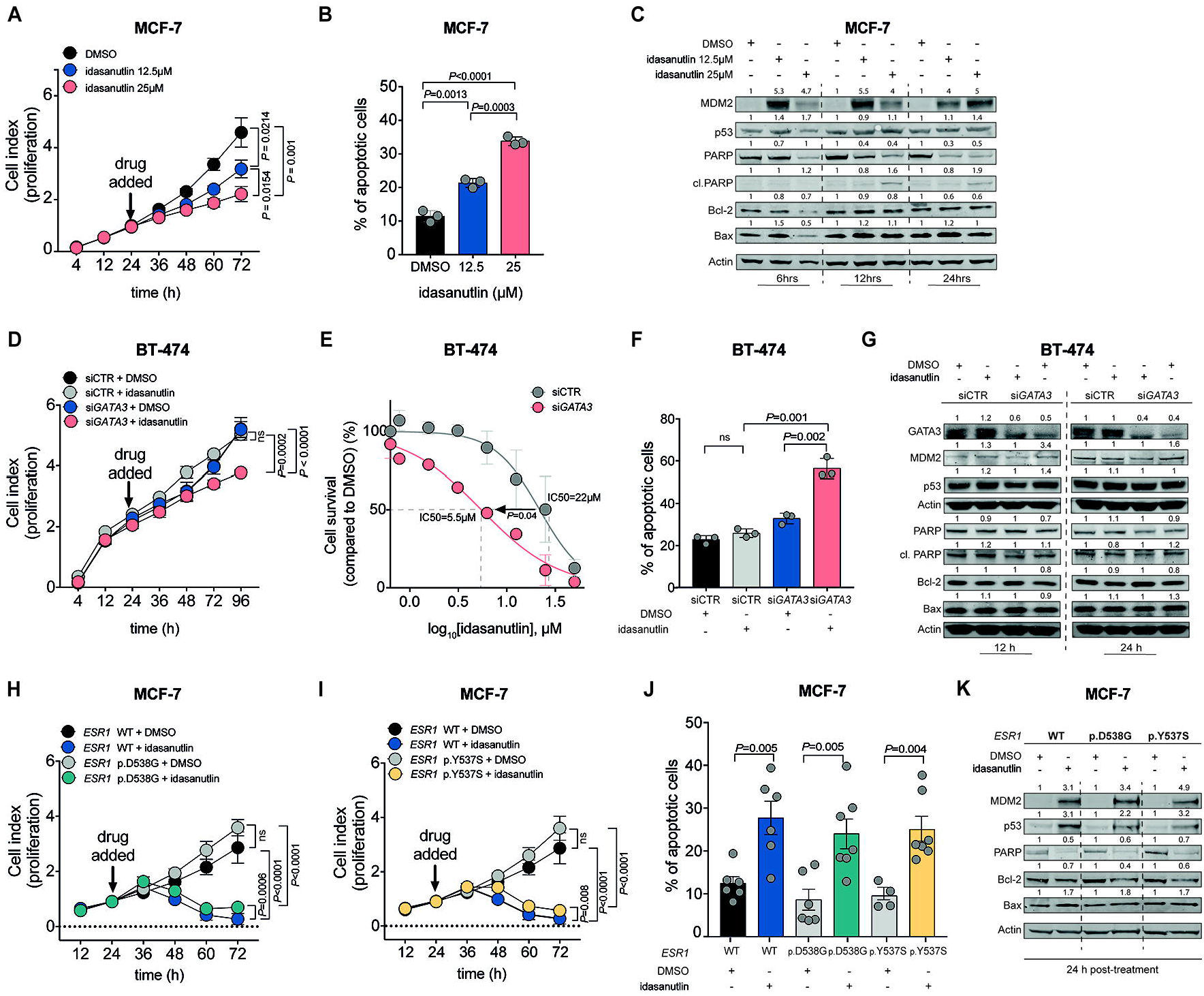
*GATA3* status determines response to MDM2 inhibitors *in vitro*. (**A,D,H,I)** Proliferation kinetics of (**A)** *GATA3*-mutant MCF-7 under increasing dosage of idasanutlin, (**D)** BT-474 upon *GATA3* silencing and/or treatment with 12.5 μM idasanutlin, (**H,I)** *GATA3*-mutant MCF-7 carrying a wild-type *ESR1* or mutant *ESR1* (p.D538G/p.Y537S) upon treatment with 12.5 μM idasanutlin (see also Figure S2A-B). (**B,F,J)** Apoptosis assay using Annexin V and propidium iodide co-staining (**B)** upon increasing dosage of idasanutlin in MCF-7, (**F)** upon *GATA3* silencing and/or treatment with 12.5 μM idasanutlin in BT-474, (**J)** upon treatment of 12.5 μM idasanutlin in MCF-7 carrying a wild-type *ESR1* or mutant *ESR1* (p.D538G/p.Y537S, see also Figure S2D). (**C,G,K)** Immunoblot showing pro- and anti-apoptotic proteins (**C)** at 6, 12 and 24 h post-treatment with DMSO, 12.5 μM and 25 μM idasanutlin in MCF-7, (**G)** at 12 and 24 h post-treatment with DMSO or 12.5 μM idasanutlin in BT-474 transfected with *GATA3* siRNA or control siRNA, (**K)** at 24 h post-treatment with DMSO or 12.5 μM idasanutlin in MCF-7 carrying wild-type or mutant *ESR1* (p.D538G/p.Y537S, see also Figure S2E). For all the western blots, quantification is relative to the loading control (actin) and normalized to the corresponding DMSO control. (**E**) Logdose response curve of idasanutlin in BT-474 transfected with *GATA3* siRNA or control siRNA (see also Figure S2C). Data are mean ± s.d. (**A,B,D,E,F,H,I,J)** n≥3 replicates. Statistical significance was determined for (**A,B,D,E,F,H,I,J)** by the two-tailed unpaired Student’s t-test.

To determine whether *GATA3* expression levels would modulate response to idasanutlin, we assessed the effect of treatment on *GATA3*-silenced BT-474 and MDA-MB134 cells. We observed that while idasanutlin treatment had no or little effect on the proliferation of the control cells, it significantly reduced cell proliferation upon *GATA3* silencing (**Figure 2D, Figure S2B**). In fact, both cell lines showed that *GATA3* silencing substantially reduced the IC50 for idasanutlin (**Figure 2E, Figure S2C)**. Flow cytometry and immunoblot further demonstrated that idasanutlin treatment induced apoptosis in both BT-474 and MDA-MB134 upon *GATA3* silencing but not in control cells (**Figure 2F-G, Figure S2D-E**).

Acquired resistance to endocrine therapy is often associated with *ESR1* activating mutations (Robinson et al., 2013) or fusion genes (Hartmaier et al., 2018). We hypothesized that *MDM2* inhibition may represent an alternative therapeutic strategy in endocrine therapy-resistant breast cancers harboring *GATA3* mutations. To test this hypothesis, we treated two derivative endocrine-resistant *GATA3*-mutant MCF-7 cell lines with knock-in *ESR1* p.D538G or p.Y537S activating mutations (Jeselsohn et al., 2018; Kuang et al., 2018) with idasanutlin. We observed that idasanutlin stopped cell proliferation in both mutant cell lines (**Figure 2H-I**). Idasanutlin also induced apoptosis and up- and down-regulated pro- and anti-apoptotic proteins, respectively (**Figure 2J-K**).

Taken together, our results demonstrate that *GATA3* loss sensitizes cells to pharmacological inhibition of MDM2 *in vitro*.

### GATA3 expression determines response to MDM2 inhibitor *in vivo*

To ascertain whether *GATA3* expression levels would also modulate response to idasanutlin *in vivo*, we performed xenotransplantation into zebrafish embryos. As a cancer model system, human cancer xenografts in zebrafish recapitulate the response to anticancer therapies of mammalian models (Fior et al., 2017; White et al., 2013). To generate the zebrafish models, we treated *GATA3*-silenced and control BT-474 cells with idasanutlin (25 μM) or vehicle (DMSO) 48 hours post-siRNA transfection. Twenty-four hours later, we labeled the cells with a red fluorescent cell tracker, injected them into the yolk sac of zebrafish embryos and screened embryos for tumor cell engraftment after four days (**Figure 3A**) (Haldi et al., 2006).

**Figure 3:**
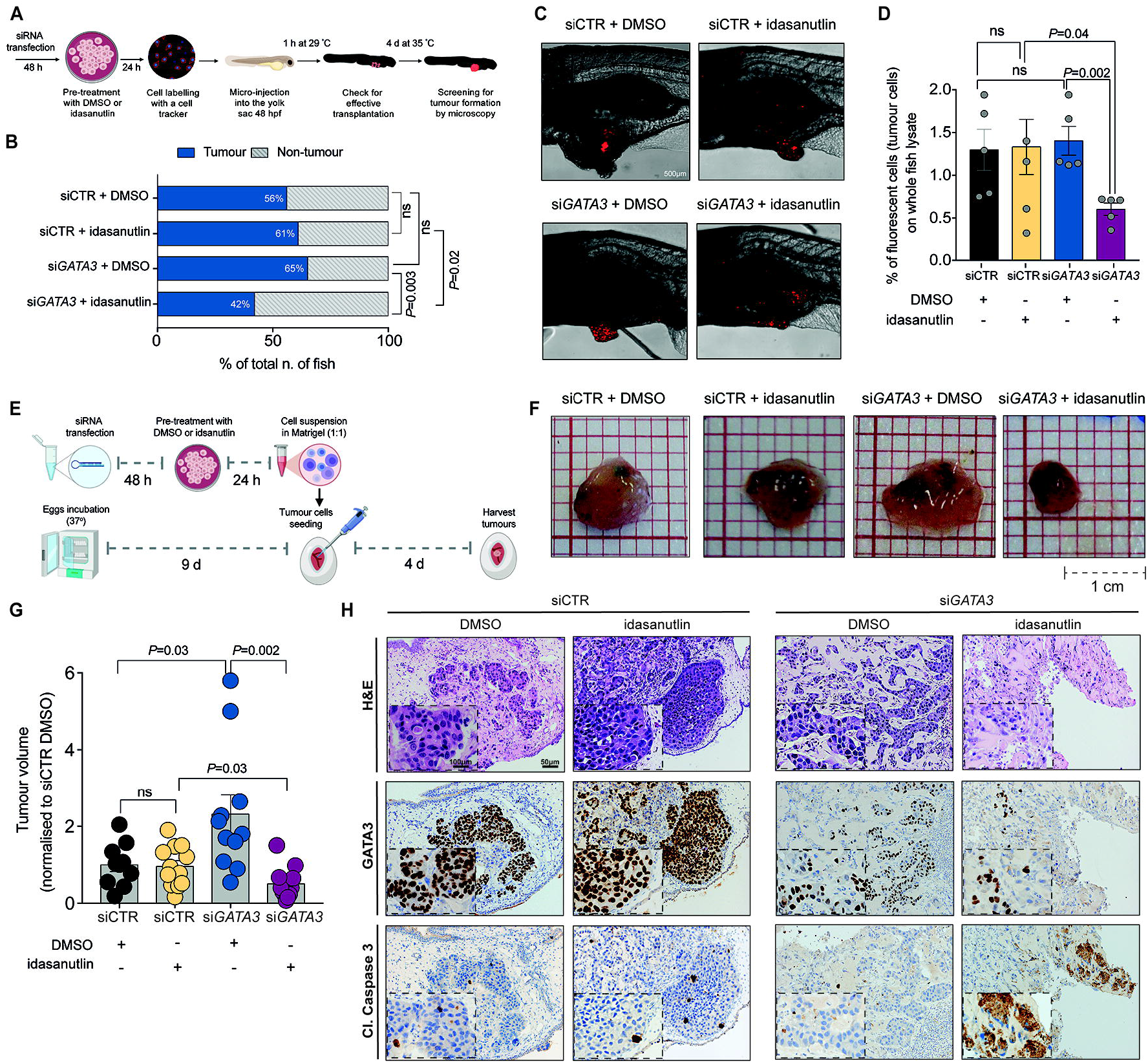
GATA3 expression determines response to MDM2 inhibitor *in vivo*. (**A)** Schematic representation of the zebrafish xenotransplantation assay. (**B)** Barplot shows the percentages of fish that harbored or did not harbor tumors upon transplantation with *GATA3*-silenced or control BT-474 cells pre-treated with idasanutlin or DMSO. In total, 70-100 embryos per group were analyzed over two independent experiments. (**C)** Representative confocal images of tumor formation in zebrafish injected with fluorescent tracker-labeled BT-474 cells with *GATA3* siRNA or control siRNA, pretreated with idasanutlin or DMSO. (**D)** FACS analysis showing the percentage of red-tracker labelled tumour cells extracted from the embryos. Error bars represent, in total, three replicates performed over two independent experiments. Each replicate represents the pooled lysate of 20-30 fish for each condition. (**E)** Schematic illustration of the CAM assay. (**F)** Photographs of *GATA3*-silenced or control BT-474 cells pre-treated with DMSO or idasanutlin implanted in CAMs and grown for four days post-implantation. (**G)** Volume of tumors derived from the CAM experiment (n≥10 tumors over three independent experiments). Values are normalized to the mean of siCTR DMSO. (**H)** Representative micrographs of BT-474 tumors extracted four days post-implantation. Tumoural cells (upper) were immunostained with GATA3 (middle) and the apoptotic marker cleaved caspase 3 (lower) in the different treatment conditions (see also Figure S3). Data are mean ± SEM (**D,G)** n≥4 replicates. Scale bars: (**C)** 500 μm, (**F)** 1cm and (**H)** 50 μm and 100 μm. Statistical significance was determined for (**B)** by two-sided Fisher’s Exact test and for (**D,G)** by the two-tailed unpaired Student’s t-test.

We observed that *GATA3*-silenced cells injected into fish were more sensitive to idasanutlin than the control (42% vs 61%, **Figure 3B**). More importantly, idasanutlin reduced tumor formation in the context of *GATA3*-silencing (42% vs 65% treated with DMSO) but not in control (61% vs 56% treated with DMSO, **Figure 3B**). Tumors derived from *GATA3*-silenced, idasanutlin-treated cells, were very small, largely consisting of small clusters of tumor cells, compared to the larger solid tumor masses derived from *GATA3*-silenced cells without idasanutlin (**Figure 3C**). To assess cell proliferation, we quantified the percentage of tumor cells present in the fish by performing FACS analysis of the fluorescence-labeled tumor cells in whole fish extracts. Consistent with the results from the tumor formation assay, idasanutlin treatment was only effective in reducing the overall percentage of tumor cells in fish injected with *GATA3*-silenced cells (purple vs DMSO-treated in blue) but not in fish injected with control (yellow vs DMSO-treated in black, **Figure 3D**), indicating that *GATA3* expression level modulates sensitivity to MDM2 inhibition *in vivo*.

The zebrafish xenograft model provides insights into the tumorigenic and proliferative capability of cancer cells. However, to assess apoptosis and to quantify tumor growth, we employed the chicken chorioallantoic membrane (CAM), a densely vascularized extraembryonic tissue, as a second *in vivo* model (Fluegen et al., 2017; Hagedorn et al., 2005). Similar to the zebrafish assay, we treated *GATA3*-silenced and control BT-474 cells with idasanutlin (25 μM) or vehicle (DMSO) for 24 hours. We then inoculated the cells into the CAMs and screened the eggs for tumour formation four days later (**Figure 3E**). In accordance with our results in the zebrafish model, idasanutlin treatment reduced the volume of tumors formed by *GATA3*-silenced cells (purple vs DMSO-treated in blue) but not in control cells (yellow vs DMSO-treated in black, **Figure 3F-G**), suggesting that *GATA3* expression modulates response to MDM2 inhibitors in the CAM model as well. We then evaluated apoptosis induction by staining tumor sections with the apoptotic marker cleaved caspase 3. Notably, only *GATA3*-silenced idasanutlin-treated tumors showed a strong positive signal for cleaved caspase 3, as well as morphological features of apoptosis (e.g. nuclear fragmentation, hypereosinophilic cytoplasm, “apoptotic bodies”, **Figure 3H, Figure S3**), demonstrating that idasanutlin induces apoptosis in the context of *GATA3* silencing *in vivo*.

Taken together, our results show that *GATA3* expression modulates response to idasanutlin in two independent *in vivo* models.

### The synthetic lethality between *GATA3* and *MDM2* is *TP53* dependent

MDM2 plays a central role in the regulation of p53 and they regulate each other in a complex regulatory feedback loop (Wu et al., 1993) (**Figure 4A**). We analyzed the frequencies of *GATA3* and *TP53* mutations in ER-positive breast cancer (Hoadley et al., 2018; Pereira et al., 2016) and observed that they are mutually exclusive (**Figure 4B**). We, therefore, hypothesized that the synthetic lethal effects between *GATA3* and *MDM2* may be p53-dependent. To test this hypothesis, we assessed cell growth and apoptosis upon single- and dual-silencing of *GATA3* and *MDM2* in the ER-positive, *GATA3*-wild-type, *TP53*-mutant (p.L194F) T-47D breast cancer cell line (**Figure S4A**). Consistent with the mutual exclusivity of *GATA3* and *TP53* mutations, *GATA3* silencing in a *TP53*-mutant context resulted in a strong reduction of cell viability and induction of apoptosis (**Figure 4C-D**). Contrary to the results obtained in cells with functional p53, *GATA3/MDM2* dual silencing did not show synthetic lethal effect (**Figure 4C-D**). If the synthetic lethal interaction between *GATA3* and *MDM2* is *TP53*-dependent, one should expect that silencing *TP53* should partially revert the phenotype. Therefore we silenced *MDM2* alone or in combination with *TP53* in the *GATA3*-mutant MCF-7 cell line (**Figure S4B**). As expected, *TP53* silencing partially rescued the effect induced by *MDM2* knock-down (**Figure 4E-F**) as well as of idasanutlin treatment (**Figure 4G-H, Figure S4C-D**) on cell growth and apoptosis, demonstrating the p53 dependency of the synthetic lethal interaction.

**Figure 4:**
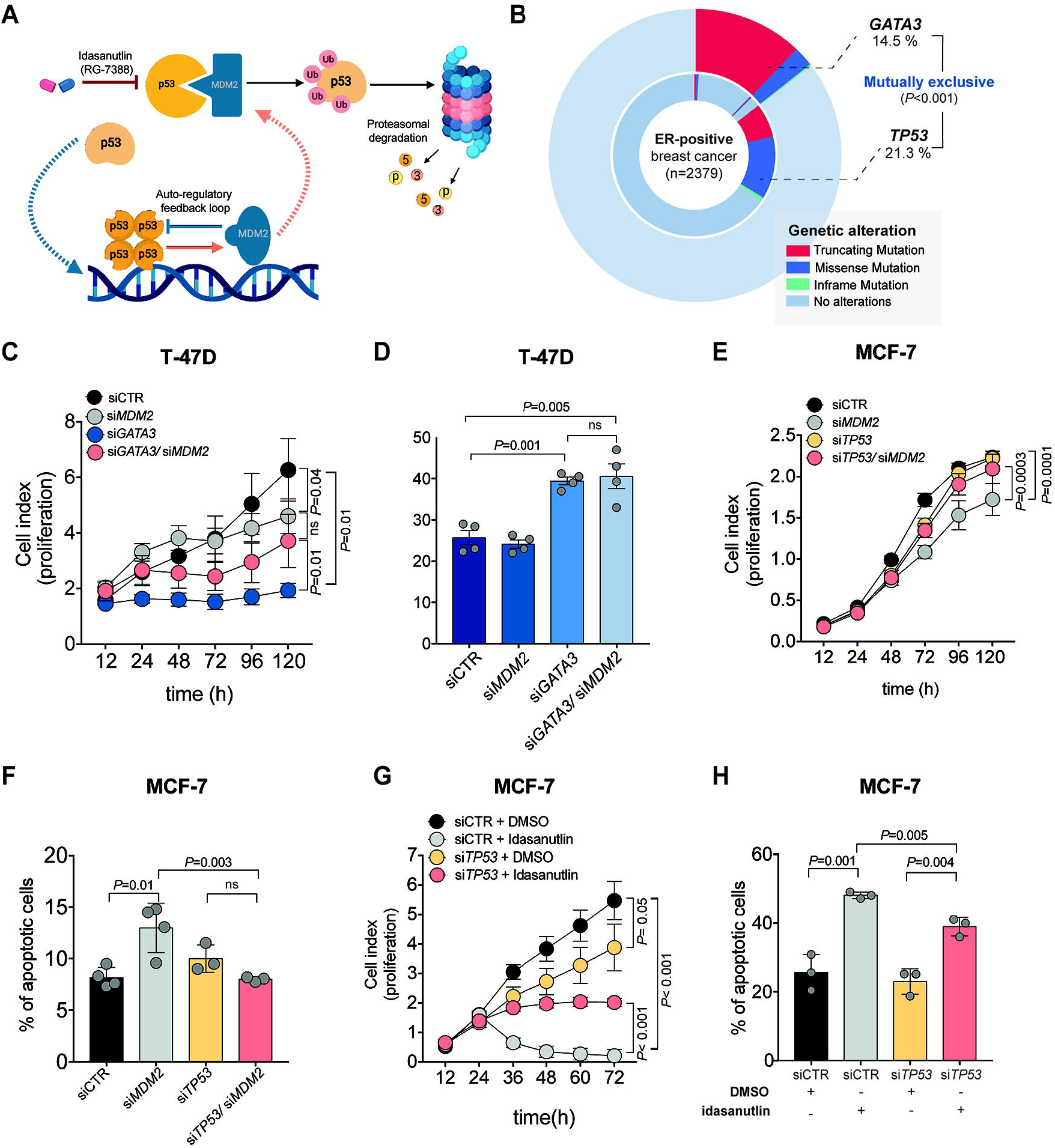
The synthetic lethality between *GATA3* and *MDM2* is *TP53* dependent. (**A)** Schematic representation on the regulatory feedback loop between MDM2 and p53. (**B**) Doughnut chart showing *GATA3* and *TP53* mutations in ER-positive breast cancer (Hoadley et al., 2018; Pereira et al., 2016). (**C**) Proliferation kinetic of *TP53*-mutant T-47D transfected with siRNA targeting *GATA3, MDM2, GATA3/MDM2*, or control (see also Figure S4A). (**D)** Percentage of apoptotic cells upon silencing of *GATA3* and *MDM2* alone or in combination in T-47D. (**E**) Proliferation kinetic of MCF-7 transfected with siRNA targeting *TP53, MDM2, TP53/MDM2*, or control (see also Figure S4B). (**F**) Percentage of apoptotic cells upon silencing of *TP53* or *MDM2* alone or in combination in MCF-7. (**G**) Proliferation kinetic of MCF-7 upon *TP53* silencing and/or treatment with 12.5 μM idasanutlin (see also Figure S4C). (**H**) Percentage of apoptotic cells upon silencing of *TP53* and/or treatment with 12.5 μM idasanutlin (see also Figure S4D). Data are mean ± s.d. (**C,D,E,F,G,H)** n≥3 replicates. Statistical significance was determined for (**B)** by one-sided Fisher’s Exact test and for (**C,D,E,F,G,H)** by the two-tailed unpaired Student’s t-test.

### *GATA3* mutational status predicts response to idasanutlin in ER-positive breast cancer patient-derived organoids

As patient-derived organoids (PDOs) have been shown to retain the molecular features of the original tumors and to better resemble tumor heterogeneity than traditional twodimensional cell culture methods derived from single cell clones, they are frequently used as *ex vivo* preclinical models for drug response prediction (Drost and Clevers, 2018; Ganesh et al., 2019; Holen et al., 2017; Vlachogiannis et al., 2018). Indeed, drug sensitivity of PDOs have been shown to mirror the patient’s response in the clinic (Ooft et al., 2019; Yao et al., 2020). We therefore tested our findings in organoids derived from three ER-positive invasive ductal breast carcinoma patients, a primary tumor and a bone metastasis harboring *GATA3* frameshift mutations (primary: p.H433fs and metastasis: p.S410fs, **Figure S5A**) and one bone metastasis carrying wild-type *GATA3* (**Figure 5A**). All tumor specimens were carrying a wild-type *TP53* gene. In accordance with the oncosuppressor role of GATA3 and our previously generated data, PDOs derived from the *GATA3*-mutant primary breast cancer significantly proliferated more compared to *GATA3* wild-type PDOs (**Figure 5B**). Importantly, all PDOs retained ERα and GATA3 expression (**Figure 5C**, **Figure S5B**). In accordance with the results generated *in vitro* and *in vivo, GATA3*-mutant PDOs showed a significant decrease in IC50 for idasanutlin (**Figure 5D, Figure S5C**). Furthermore, viability assay revealed that both *GATA3*-mutant PDOs treated with idasanutlin significantly proliferated less (~50% for the primary and ~25% for the bone metastasis) compared to the wild-type PDOs (**Figure 5E-F, Figure S5D**). In particular, upon treatment with 1.5 μM idasanutlin *GATA3* wild-type PDOs were still 100% alive compared to their DMSO control counterparts while only 40% (primary) and 60% (metastasis) *GATA3*-mutant PDOs survived (**Figure 5E-F, Figure S5C**).

**Figure 5:**
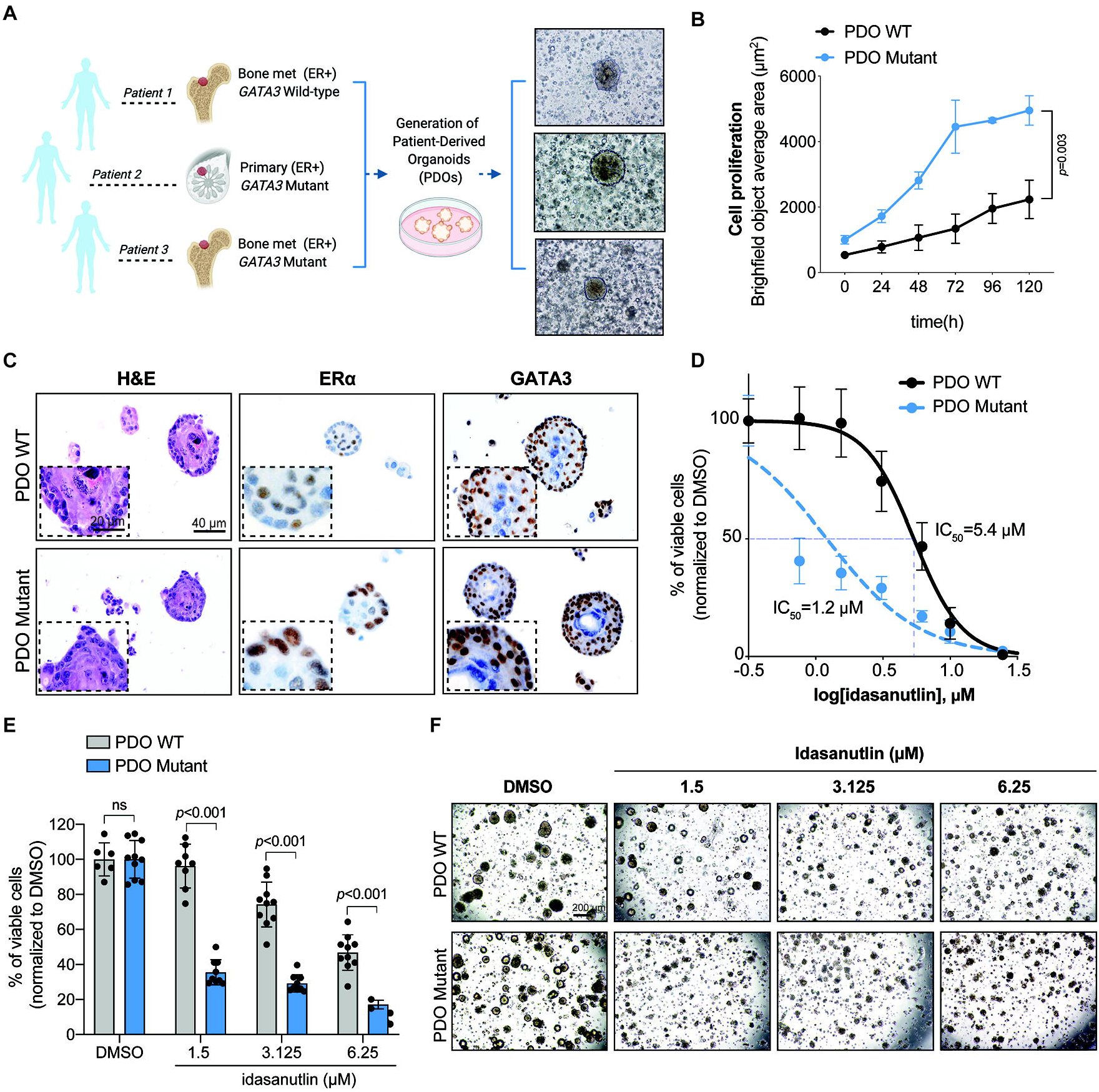
*GATA3* mutations predict response to idasanutlin in patient-derived organoids. (**A)** Schematic representation and representative microscopy pictures of the generation of organoids from n=3 ER-positive breast cancers (see also Figure S5A). (**B**) Proliferation kinetics of *GATA3*-wild-type PDOs (black, patient 1) and *GATA3*-mutant PDOs (blue, patient 2). (**C**) Representative micrographs of H&E, ER□ and GATA3 immuno-staining on the PDOs (see also Figure S5B). (**D**) Log-dose response curve of idasanutlin in *GATA3*-wild-type (IC50=5.4 μM) or *GATA3*-mutant (1.2 μM) PDOs (see also Figure S5C). (**E)** Percentage of viable cells upon treatment with different dosages of idasanutlin in *GATA*3-wild-type (gray) or *GATA*3-mutant (blue) PDOs (see also Figure S5D). (**F**) Representative micrographs of PDOs after five days of treatment with different dosages of idasanutlin. Scale bars are 20 and 40 μm for (**C**) and 200 μm for (**F**). Data are mean ± SD, n≥4 replicates from two independent experiments (**D, E**). Statistical significance was determined for (**B,E)** by two-tailed unpaired Student’s t-tests.

Our *ex vivo* data further support the use of MDM2 inhibition in the treatment of *GATA3*-mutant ER-positive breast cancer patients.

### The synthetic lethality between *GATA3* and *MDM2* acts via the PI3K-Akt-mTOR signaling pathway

To investigate the putative mechanisms driving the synthetic lethality, we analyzed the gene expression changes induced by concurrent *GATA3* loss and *MDM2* silencing. RNA-sequencing analysis of the *MDM2*-silenced MCF-7 cells and dual *GATA3/MDM2-silenced* MDA-MB134 cells revealed 20 commonly dysregulated pathways (**Figure 6A**). As expected, pathways related to p53 and apoptosis were significantly up-regulated in both cell lines, while many proliferation-related pathways such as *E2F* and *MYC* targets were down-regulated (**Figure 6B**). Interestingly, the mTORC1 signaling pathway was among the most significantly down-regulated pathways in both cell lines. Indeed, we confirmed that *MDM2* silencing in the *GATA3*-mutant MCF7 cells (**Figure S6A**) reduced phospho-Akt, phospho-S6, as well as phospho-GSK3β, compared to control cells (**Figure 6C**), indicating the downregulation of the mTOR pathway. Similarly, in BT-474 cells, dual *GATA3/MDM2* silencing reduced levels of phospho-Akt, phospho-S6 and phospho-GSK3β and induced apoptosis (**Figure 6D and Figure S6A**). By contrast, phospho-Akt levels were higher when only *GATA3* was silenced (**Figure 6D and Figure S6B**). Pharmacological inhibition of MDM2 in *GATA3*-silenced BT-474 cells also resulted in a reduction in phospho-Akt, phospho-S6 and phospho-GSK3β (**Figure S6B**). To determine whether deregulation of the mTOR signaling cascade could also be observed *in vivo*, we stained the tumors in our CAM model with phospho-S6 and phospho-Akt. Indeed, in tumors derived from *GATA3*-silenced BT-474 cells, both phospho-S6 and phospho-Akt were drastically reduced upon treatment with idasanutlin, while in tumors derived from control cells, idasanutlin treatment did not have an effect on mTOR signaling (**Figure 6E**, **Figure S6C**).

**Figure 6:**
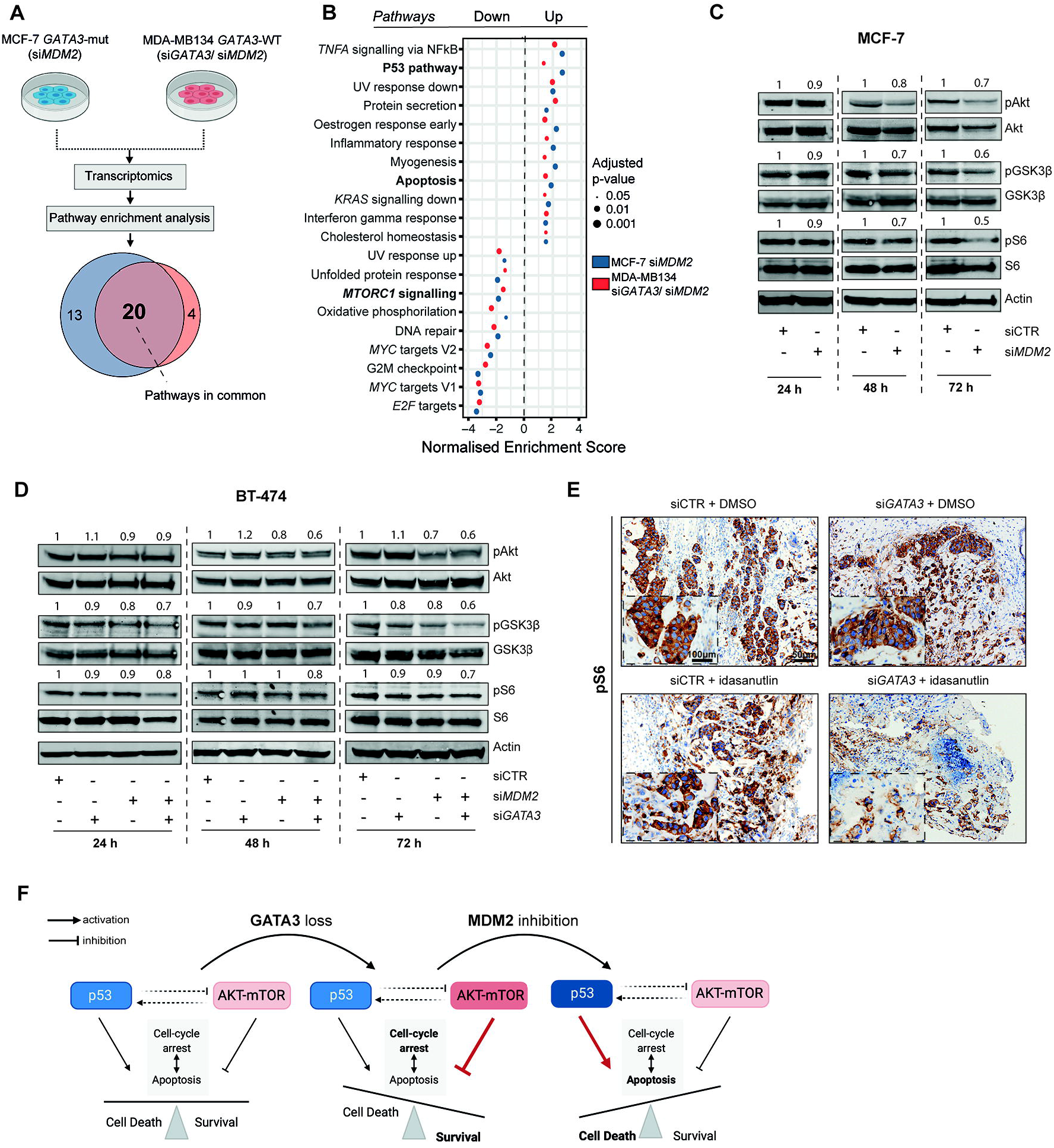
The synthetic lethality between *GATA3* and *MDM2* acts via the PI3K-Akt-mTOR signaling pathway. (**A)** Schematic representation of the RNA-seq experimental setup to identify gene expression changes induced by concurrent *GATA3* loss and *MDM2* inhibition. Venn diagram shows the number of pathways enriched in both MCF-7 with *MDM2* siRNA and MDA-MB134 with *GATA3* siRNA and *MDM2* siRNA. (**B)** Normalized enrichment scores of significantly up- and down-regulated pathways identified by gene set enrichment analysis in both MCF-7 and MDA-MB134. Size of the dots is proportional to the adjusted p-value as indicated in the legend. (**C,D)** Immunoblot showing markers of mTOR signaling pathway activation at 24, 48 and 72 hours post-siRNA transfection in (**C)** MCF-7 cells upon *MDM2* silencing and (**D)** BT-474 cells upon *GATA3* and/or *MDM2* silencing (see also Figure S6A). For all the western blots, quantification is relative to the loading control (actin) and normalized to the corresponding siCTR. (**E)** Representative immunohistochemistry micrographs of phospho-S6 stainings in BT-474 tumors extracted four days postimplantation in the CAM model (see also Figure S6C). (**F)** Schematic representation of the mechanistic hypothesis. Scale bars: (**E)** 50 μm and 100 μm. Statistical significance was determined for (**B)** by *fgsea*.

We, therefore, hypothesized that GATA3 loss may induce addiction to mTOR signaling in breast cancer cells. In support of our hypothesis, we observed that, in ER-positive breast cancers, genetic alterations in *GATA3* are significantly mutually exclusive with those in both *PI3KCA* and *PTEN* (**Figure S6D-E**). Furthermore, differential gene expression and pathway enrichment analyses between *GATA3*-mutant and *GATA3*-wild type ER-positive breast cancers and between ER-positive breast cancers with low and high *GATA3* expression levels also showed significant enrichment for the mTORC1 signaling pathway (**Figure S6F-G**).

Taken together, our results show that the synthetic lethality between GATA3 and MDM2 acts via the PI3K-Akt-mTOR signaling pathway.

## Discussion

*GATA3* is mutated in 12-18% of breast cancer (Bertucci et al., 2019; Hoadley et al., 2018) with predominantly frameshift mutations and loss of GATA3 expression is strongly associated with failure to respond to hormonal therapy and poor prognosis (Gulbahce et al., 2013). Here we describe a novel synthetic lethal interaction between *GATA3* and *MDM2* in ER-positive breast cancer. In particular, we showed that, in the context of truncating *GATA3* mutation, inhibition of MDM2 hampers cell proliferation and induces apoptosis *in vitro*, in two independent *in vivo* models and in two *GATA3*-mutant PDOs. In the context of wild-type *GATA3*, the same effect was achieved by dual *GATA3* and *MDM2* inhibition, regardless of *HER2* status. We further showed that *GATA3* expression level modulates response to MDM2 inhibitor. Our results thus support MDM2 as a therapeutic target in the substantial fraction of ER-positive, *GATA3*-deficient breast cancer. Of note, although *TP53* mutations are a major driver of resistance to MDM2 inhibitors (Khoo et al., 2014; Marcellino et al., 2019), very few *GATA3*-mutant ER-positive breast cancers harbor *TP53* mutations. Thus the presence of *TP53* mutations is not expected to preclude the use of MDM2 inhibitors in the vast majority of these patients. With MDM2 inhibitors, such as idasanutlin, widely available, our findings allow the rational design of clinical trials to evaluate the in-patient efficacy of MDM2 inhibitors and to specifically evaluate GATA3 status as a predictive biomarker of response. Given that GATA3 loss of expression has also been associated with poor prognosis in other cancer types (Miyamoto et al., 2012; Nguyen et al., 2013), we expect our finding to have far-reaching implications beyond ER-positive breast cancer.

We showed that the synthetic lethality between *GATA3* and *MDM2* is p53-dependent and acts via the PI3K/Akt/mTOR pathway. It is well known that in normal conditions p53 and the PI3K/Akt/mTOR pathway co-regulate cell cycle arrest and apoptosis leading to homeostasis between cell death and survival (Feng et al., 2005; Vousden and Lu, 2002). Our results suggest that in breast cancer cells, *GATA3* loss-of-function (via genetic alterations or other mechanisms) activates the PI3K/Akt/mTOR pathway and leads to resistance to apoptosis. In this context, MDM2 inhibition, with consequent p53 up-regulation and mTOR signaling downregulation, pushes the cells toward cell death (**Figure 6F**). In support of this model, downregulation of *GATA3* has been directly linked to Akt kinase activation in breast and prostate cancers (Nguyen et al., 2013; Werner et al., 2015; Yu et al., 2019). It has also been reported that upon adaptation to hormone deprivation, breast cancer cells rely heavily on PI3K signaling and that inhibition of PI3K and mTOR induces apoptosis in these cells (Miller et al., 2010). Furthermore, our model is also supported by the observed synergistic effect of dual MDM2 and PI3K/Akt/mTOR inhibition (Kojima et al., 2008; Moreno-Smith et al., 2017). Our hypothesis, however, may only partially explain the synthetic lethality between GATA3 and MDM2. Further studies are required to fully dissect the mechanism of action.

Our findings have important clinical implications for several subsets of ER-positive breast cancer. First, our findings suggest that *GATA3* mutations may drive resistance to hormonal therapy by upregulating the PI3K/Akt/mTOR pathway, suggesting that inhibition of MDM2 may help overcome the resistance to PI3K/Akt/mTOR inhibition and/or to endocrine therapy in *GATA3*-deficient breast cancer. Second, the mutual exclusivity between genetic alterations in *GATA3* and genes in the PI3K pathway suggests the PI3K inhibitors may be effective in the context of *GATA3* mutations. It would be clinically relevant to test if the synergistic effect of dual MDM2 and PI3K/Akt/mTOR inhibition is even stronger in the context of *GATA3* mutation. Third, given that aberrant activation of the PI3K pathway has been implicated in resistance to HER2-targeted therapy (Nagata et al., 2004), one might hypothesize that *GATA3* mutations may be a mechanism of resistance to HER2-targeted therapy and that MDM2 inhibitors may act synergistically with trastuzumab in ER+/HER2+, *GATA3*-mutant breast cancers.

Despite the profound therapeutic implications, the synthetic lethality between GATA3 and MDM2 had never been reported. This unexpected finding was the result of the availability of large-scale, unbiased screening of genetic interactions in a large panel of cell lines (McDonald et al., 2017) as well as a statistical algorithm (Srivatsa et al.) powerful enough to detect such interaction even when the number of cell lines harboring *GATA3* mutation is small (n=2). Our study exemplifies how perturbation screens can lead to pre-clinical hypotheses that can be rapidly tested and translated into therapeutic candidates.

## Supporting information

Key Resources Table

## Acknowledgments

We would like to thank Stefano Medagli who helped with the generation of the figures. We would like to thank Dr. Rachael Natrajan for sharing breast cancer cell lines. RNA-sequencing analysis was performed at sciCORE scientific computing center at the University of Basel.

## Author contributions

C.K.Y.N. and S.P. conceived and supervised the study, interpreted the results, wrote and edited the manuscript with G.B. and M.C.-L.; G.B., C.K.Y.N. and S.P. made final edits, figures and completed the paper; G.B., M.C.-L., S.T.-M., and V.P. performed molecular experiments and G.B. analysed the data; M.K., M.D.M, M.K.-D.J., and C.L. provided the technical expertise for the *in vivo* experiments performed by G.B., M.C.-L, S.T.-M and M.K.; G.B. and F.P. performed the *ex vivo* experiments; S.S., H.M. and N.B. developed the statistical model for the analysis of the DRIVE data; C.E. and L.M.T. analyzed the histological stains and provided pathology expertise; J.G. and V.K. performed bioinformatic analysis of the RNA-sequencing and the publicly available data; R.M.J. provided the *ESR1* mutant models and provided critical input in the discussion of the results. F.-C.B. provided critical input in the discussion of the results; A.D, E.Mo, E.Ma. provided the human *GATA3* wild-type and mutant tumor models and critical input in the discussion of the results. All authors agreed to the final version of the manuscript.

## STAR Methods

### Cell lines

ER-positive breast cancer cell lines MCF-7 (*GATA3*-mutant p.D335Gfs; *TP53* wild-type), BT-474 (*GATA3* wild-type, *TP53*-mutant p.E285K with retained transactivation activity (Jordan et al., 2010)), MDA-MB134 (*GATA3* wild-type; *TP53* wild-type) and T-47D (*GATA3* wild-type, *TP53* mutant p.L194F) were kindly provided by Dr. Rachael Natrajan from The Institute of Cancer Research (London, UK), authenticated by short tandem repeat profiling. MCF-7 cell lines with knock-in mutations in the *ESR1* gene (p.Y537S and p.D538G) were provided by Dr. Jeselsohn (Jeselsohn et al., 2018). All cell lines were monitored regularly for mycoplasma contamination by PCR using specific primers as described previously (Uphoff and Drexler, 2011). All cell lines were maintained under the condition as recommended by the provider. Briefly, all cell lines were cultured in DMEM supplemented with 5% Fetal Bovine Serum, non-essential amino-acids and antibiotics (Penicillin/Streptomycin). The cells were incubated at 37°C in a humidified atmosphere containing 5% CO2. Exponentially growing cells were used for all *in vitro* and *in vivo* studies.

### Gene knockdown by siRNAs

Transient gene knockdown was conducted using ON-TARGET plus siRNA transfection. ON-TARGET plus SMARTpool siRNAs against human *GATA3, MDM2, TP53*, ON-TARGET plus SMARTpool non-targeting control and DharmaFECT transfection reagent were all purchased from GE Dharmacon (**Key Resources Table)**. Transfection was performed according to the manufacturer’s protocol. Briefly, log-phase ER-positive breast cancer cells were seeded at approximately 60% confluence. Because residual serum affects the knockdown efficiency of ON-TARGET plus siRNAs, growth medium was removed as much as possible and replaced by serum-free medium (Opti-MEM). siRNAs were added to a final concentration of 25 nM, unless otherwise specified (Note: siRNAs targeting different genes can be multiplexed). Cells were incubated at 37°C in 5% CO2 for 24, 48 and 72 h (for mRNA analysis) or for 48 and 72 h (for protein analysis). To avoid cytotoxicity, the transfection medium was replaced with a complete medium after 24 h.

### RNA extraction and relative expression by qRT-PCR

Total RNA was extracted from cells at 75% confluence using TRIZOL (**Key Resources Table)** according to the manufacturer’s guidelines. cDNA was synthesized from 1 μg of total RNA using SuperScript™ VILO™ cDNA Synthesis Kit. All reverse transcriptase reactions, including no-template controls, were run on an Applied Biosystem 7900HT thermocycler. The expression for all the genes was assessed using SYBR and all qPCR experiments were conducted at 50°C for 2min, 95°C for 10min, and then 40 cycles of 95°C for 15 s and 60°C for 1min on a QuantStudio 3 Real-Time PCR System (Applied Biosystems). The specificity of the reactions was verified by melting curve analysis. Measurements were normalized using *GAPDH* level as reference. The fold change in gene expression was calculated using the standard ΔΔCt method (Livak and Schmittgen, 2001). All samples were analyzed in triplicate. List of primers is available in **Key Resources Table**.

### Immunoblot

Total proteins were extracted by directly lysing the cells in Co-IP lysis buffer (100mmol/L NaCl, 50mmol/L Tris pH 7.5, 1mmol/L EDTA, 0.1% Triton X-100) supplemented with 1x protease inhibitors and 1x phosphatase inhibitors. Cell lysates were then treated with 1x reducing agent, 1x loading buffer, boiled and loaded onto neutral pH, pre-cast, discontinuous SDS-PAGE mini-gel system. After electrophoresis, proteins were transferred to nitrocellulose membranes using the Trans-Blot Turbo Transfer System (Bio-Rad). The transblotted membranes were blocked for 1 h in TBST 5% milk and then probed with appropriate primary antibodies (from 1:200 to 1:1000) overnight at 4°C. List of antibodies and working concentrations are available in **Key Resources Table**. Next, the membranes were incubated for 1 h at room temperature with fluorescent secondary goat anti-mouse (IRDye 680) or antirabbit (IRDye 800) antibodies (both from LI-COR Biosciences). Blots were scanned using the Odyssey Infrared Imaging System (LI-COR Biosciences) and band intensity was quantified using ImageJ software. The ratio of proteins of interest/loading control in idasanutlin-treated samples were normalized to their DMSO-treated control counterparts. All experiments were performed and analyzed in triplicate.

### Drug treatment

10×10^3^ exponentially growing cells were plated in a 96-well plate. After 24 h, cells were treated with serial dilution of RG7388-idasanutlin (**Key Resources Table)** or dimethyl sulfoxide (DMSO). DMSO served as the drug vehicle, and its final concentration was no more than 0.1%. Cell viability was measured after 72 h using CellTiter-Glo Luminescent Cell Viability Assay reagent. Results were normalized to the vehicle (DMSO).

For the treatment experiments of the PDOs, PDOs were plated as single cells in a 96-well plate at a density of 1 × 10^4^ cells in 10 μl Matrigel droplets. Prior to treatment, cells were allowed to recover and form organoids for 2 days. At day 3, idasanutlin at different dilutions (2 fold, from 25 to 1.5 μM) was added to the medium, and cell viability was assessed after 5 days using CellTiter-Glo 3D reagent (**Key Resources Table)**. Luminescence was measured on Varioskan Microplate Reader (ThermoFisher Scientific). Results were normalized to DMSO control. All experiments were performed in triplicate. Results are shown as mean ± SD. Curve fitting was performed using Prism (GraphPad) software and the nonlinear regression equation.

### Proliferation assay

For cell lines cell proliferation was assayed using the xCELLigence system (RTCA, ACEA Biosciences) as previously described (Andreozzi et al., 2016). Background impedance of the xCELLigence system was measured for 12 s using 50 μl of room temperature cell culture media in each well of E-plate 16. Cells were grown and expanded in tissue culture flasks as previously described (Andreozzi et al., 2016). After reaching 75% confluence, cells were washed with PBS and detached from the flasks using a short treatment with trypsin/EDTA. 5,000 cells were dispensed into each well of an E-plate 16. Cell growth and proliferation were monitored every 15 min up to 120 hours via the incorporated sensor electrode arrays of the xCELLigence system, using the RTCA-integrated software according to the manufacturer’s parameters. In the case of transient siRNA transfection, cells were detached and plated on xCELLigence 24 h post-transfection. For all the experiments with idasanutlin (RG7388), the drug or DMSO were added to the cells 24 h post-seeding on the xCELLigence system, as indicated on the figures. All experiments were performed in triplicate. Results are shown as mean ± SD.

For the PDOs, cell proliferation was assayed using the Incucyte S3 Live-Cell analysis system (Sartorius). Briefly, PDOs were plated as single cells in a 96-well plate at a density of 1 × 10^4^ cells in 10 μl Matrigel droplets and allowed to recover and form organoids for 2 days. At day 3 after seeding, the 96-well plate was placed in the Incucyte incubator where a camera automatically acquired images of each well every 8 hours (up to 120 hours). Kinetic curves were obtained using the Incucyte analysis software and the Spheroid analysis module. Cell proliferation was calculated as brightfield object average area. All experiments were performed in duplicate. Results are shown as mean ± SD.

### Apoptosis analysis by flow cytometry

Cells were collected 72 hours post-siRNA transfection and 48 hours post-treatment with idasanutlin (RG7388) respectively, stained with annexin V (AnnV) and propidium iodide (PI), and analysed by flow cytometry using the BD FACSCanto II cytometer (BD Biosciences, USA). Briefly, cells were harvested after incubation period and washed twice by centrifugation (1,200g, 5min) in cold phosphate-buffered saline (DPBS; Gibco, CO; #14040133). After washing, cells were resuspended in 0.1ml AnnV binding buffer 1X containing fluorochrome-conjugated AnnV and PI (PI to a final concentration of 1 μg/ml) and incubated in darkness at room temperature for 15min. As soon as possible cells were analyzed by flow cytometry, measuring the fluorescence emission at 530 nm and >575 nm. Data were analyzed by FlowJo software version 10.5.3.

### Zebrafish xenografts

Animal experiments and zebrafish husbandry were approved by the “Kantonales Veterinaeramt Basel-Stadt” (haltenewilligung: 1024H) in Switzerland and the experiments were carried out in compliance of ethics regulation. Zebrafish were bred and maintained as described previously (Nusslein-Volhard and Dahm, 2002). Staging was done by hours postfertilization (hpf) as described previously (Kimmel et al., 1990) and according to FELASA and Swiss federal law guidelines. Zebrafish wild-type Tuebingen strains were used in this study. 48 h post-siRNA transfection, *GATA3*-silenced and control BT-474 cells were treated for 24 h with idasanutlin (25 μM). After harvesting, cells were labeled with a lipophilic red fluorescent dye (CellTracker™ CM-DiI), according to the manufacturer’s instructions. Zebrafish were maintained, collected, grown and staged in E3 medium at 28.5°C according to standard protocols(Choi et al., 2007). For xenotransplantation experiments, zebrafish embryos were anesthetized in 0.4% tricaine at 48 h (hpf) and 200 *GATA3*-silenced or control BT-474 cells were micro-injected into the vessel-free area of the yolk sac. After injection, embryos were incubated for 1 hour at 28.5–29°C for recovery and cell transfer verified by fluorescence microscopy. Embryos were examined for the presence of a fluorescent cell mass localized at the injection site in the yolk sac or hindbrain ventricle. Fish harboring red cells were incubated at 35°C as described previously (Haldi et al., 2006; Konantz et al., 2012). On assay day 4, embryos were screened by fluorescence microscopy for (a) normal morphology, (b) a visible cell mass in the yolk or hindbrain ventricle, using a Zeiss SteREO Discovery V20 microscope and the number of tumor-bearing fish quantified. The screening was performed independently by two scientists. For each condition, 70 to 100 fish were analyzed over two experiments. Representative pictures were taken using a Nikon CSU-W1 spinning disk microscope. To assess cell proliferation, fish were furthermore dissociated into single cells as described previously (Carapito et al., 2017; Svoboda et al., 2016) and the number of fluorescence-labeled cells was then determined using flow cytometry on a BD FACSCanto II cytometer for CM-DiI–positive cells. For each condition, 20 to 30 fish were analyzed. Each experiment was repeated twice.

### Chorioallantoic membrane (CAM)

Fertilized chicken eggs were obtained from Gepro Geflügelzucht AG at day 1 of gestation and were maintained at 37°C in a humidified (60%) incubator for nine days (Zijlstra et al., 2002) At this time, an artificial air sac was formed using the following procedure: a small hole was drilled through the eggshell into the air sac and a second hole near the allantoic vein that penetrates the eggshell membrane. A mild vacuum was applied to the hole over the air sac in order to drop the CAM. Subsequently, a square 1cm window encompassing the hole near the allantoic vein was cut to expose the underlying CAM (Zijlstra et al., 2002). After the artificial air sac was formed, BT-474 cells growing in tissue culture were inoculated on CAMs at 2×10^6^ cells per CAM, on three to four CAMs each. Specifically, 48 h post-siRNA transfection, *GATA3*-silenced and control BT-474 cells were treated with idasanutlin (25 μM). 24 hours post-treatment, cells were detached from the culture dish with Trypsin, counted, suspended in 20 μl of medium (DMEM) and mixed with an equal volume of Matrigel. To prevent leaking and spreading of cells, a 8mm (inner diameter) sterile teflon ring (removed from 1.8 ml freezing vials, Nunc, Denmark) was placed on the CAMs and the final mixture was grafted onto the chorioallantoic membranes inoculating the cells with a pipette inside the ring (Kim et al., 1998). Embryos were maintained at 37 °C for 4 days after which tumors at the site of inoculation were excised using surgical forceps. Images of each tumor were acquired with a Canon EOS 1100D digital camera. Surface measurements were performed by averaging the volume (height*width*width) of each tumor using ImageJ, as previously described (Lauzier et al., 2019). Total n≥10 tumors for each condition were analyzed over three independent experiments.

### Patient-derived organoids (PDOs) generation and sequencing

PDOs were derived from ER-positive breast cancer patient-derived xenografts (PDX) generated at Institut Curie (Paris, France). The two metastasis-derived PDX (patients 1 and 3 in Figure 5A) correspond to PDX HBCx-139 (*GATA3*-wild-type) and HBCx-137 (*GATA3*-mutant) were established from spinal metastases of patients progressing on endocrine treatments, as previously described (Montaudon et al., 2020). The second *GATA3*-mutant PDX (HBCx-169) was established from the surgical specimen obtained at mastectomy from a *de novo* stage IV breast cancer patient.

Briefly, upon removal from the donor, mouse tissue was placed in MACS Tissue Storage Solution (**Key Resources Table)** and immediately shipped overnight on ice. Upon arrival, the tissue was immediately processed to generate PDO as previously described (Sachs et al., 2018). Briefly, the tissue was cut into small pieces and digested in 5 mL advanced DMEM/F-12, containing collagenase IV, DNase IV, hyaluronidase V, BSA and LY27632 (**Key Resources Table)** for 1 h and 30 mins at 37°C under slow rotation and vigorous pipetting every 15 mins. The tissue lysate was then filtered through a 100 μM cell strainer, centrifuged at 300 g for 10 mins and then treated with Accutase (**Key Resources Table)** for 10 mins at room temperature to dissociate the remaining fragments. After 5 mins of centrifugation at 300 g, the cell pellet was finally suspended with growth factor reduced Matrigel (**Key Resources Table)** and seeded as drops in a tissue-culture dish. After polymerization of Matrigel, medium supplemented with growth factors (Sachs et al., 2018) was added to the cells. Medium was changed every 3 days and organoids were passaged after dissociation with 0.25% Trypsin-EDTA (**Key Resources Table**).

DNA was extracted from snap frozen organoids pellets using the DNeasy Blood & Tissue Kit (Qiagen, Cat No./ID: 69504). Sanger sequencing was performed as previously described (Weinreb et al., 2014) using custom designed primers (**Key Resources Table**).

### Immunohistochemistry

Tumors were fixed in 10% Paraformaldehyde (PFA) immediately after excision from the CAM. PFA-fixed and paraffin-embedded tumors were cut as 3.5 μm thick sections. Hematoxylin and eosin (H&E) staining was performed according to standard protocols. Tissue sections were rehydrated and immunohistochemical staining was performed on a BOND-MAX immunohistochemistry robot (Leica Biosystems) with BOND polymer refine detection solution for DAB, using anti-GATA3, cleaved caspase 3, phospho-Akt or phospho-S6 (**Key Resources Table)** primary antibodies as substrate. Estrogen receptor immunostain was performed as described previously (Soysal et al., 2015). Photomicrographs of the tumors were acquired using an Olympus BX46 microscope. All stained sections were evaluated blindly by two independent pathologists.

### RNA sequencing and pathway analysis

Biological duplicates were generated for all the samples analyzed. Total RNA was extracted from cells at 75% confluence using TRIZOL (**Key Resources Table)** according to the manufacturer’s guidelines. RNA samples were treated with Turbo DNase (AM 1907, Thermo Fisher Scientific) and quantified using a Qubit Fluorometer (Life Technologies). RNA integrity was measured using the Agilent Bioanalyzer 2100 (Agilent Technologies).

Library generation was performed using the TruSeq Stranded mRNA protocol (Illumina). Paired-end RNA sequencing was performed on the Illumina NovaSeq 6000 platform using the 2×100bp protocol according to the manufacturer’s guidelines. Reads were aligned to the GRCh37 human reference genome using STAR 2.7.1 (Dobin et al., 2013), and transcript quantification was performed using RSEM 1.3.2 (Li and Dewey, 2011). Genes without at least ten assigned reads in at least two samples were discarded. Counts were normalized using the median of ratios method from the DESeq2 package (Love et al., 2014) in R version 3.6.1 (https://www.R-project.org/). Differential expression analysis was performed using the DESeq2 Wald test. Gene set enrichment analysis was performed using the *fgsea* R package (Sergushichev) and the Hallmark gene set from the Molecular Signatures Database (Liberzon et al., 2015), using the ranked t statistics from the DESeq2 Wald test. Pathways with false discovery rate (FDR) < 0.05 were considered to be significant. Results were visualized using ggplot2 (Wickham, 2009).

### Analysis of The Cancer Genome Atlas (TCGA) data

ER-positive breast cancer mutation annotation file for variant calling pipeline mutect2, FPKM gene expression data and raw read counts of the TCGA BRCA project were downloaded using *TCGAbiolinks (Colaprico et al., 2016)* package. Tumor samples were classified as *GATA3*-mutant (n=122) and *GATA3*-wild type (n=596) according to the *GATA3* mutation status. Samples with *GATA3* mRNA expression in the bottom and top quartile were classified as *GATA3*-low (n=200) and *GATA3*-high (n=204), respectively. *edgeR* package (Robinson et al., 2010) was used for differential expression analysis and the genes with low expression (<1 log-counts per million in ≥ 30 samples) were filtered out. Normalization was performed using the “TMM” (weighted trimmed mean) method (Robinson and Oshlack, 2010) and differential expression was assessed using the quasi-likelihood F-test. Gene set enrichment analysis of all analysed genes ranked based on signed p-value according to the direction of the log-fold change was performed using the *fgsea* package (Sergushichev). Hallmark gene sets from Molecular Signatures Database (Liberzon et al., 2015) were used to identify significantly up-/down-regulated pathways. Pathways with FDR < 0.05 were considered significantly regulated.

### Analysis of mutual exclusive genetic alterations

ER-positive breast cancer mutational data for the *GATA3, TP53, PIK3CA* and *PTEN* genes and copy number status for *PTEN* derived from the TCGA PanCancer Atlas(Hoadley et al., 2018) and the METABRIC dataset (Pereira et al., 2016) were obtained using cBioportal (Cerami et al., 2012). A total of 2379 samples were used for the analysis. Mutual exclusivity of somatic mutations in *GATA3, TP53, PIK3CA* and *PTEN* and deep deletions for *PTEN* were calculated using one-sided Fisher’s Exact and *P*<0.05 was considered statistically significant.

### Quantification and statistical analysis

Statistical analyses were conducted using Prism software v7.0 (GraphPad Software, La Jolla, CA, USA). For *in vitro* studies, statistical significance was determined by the two-tailed unpaired Student’s t-test. For comparison involving multiple time points, statistical significance was determined by the two-tailed unpaired multiple Student’s t-test. A *P* value < 0.05 was considered statistically significant. For all figures, ns, not significant. For *in vivo* studies two-sided Fisher’s Exact was used to compare the number of tumor-harboring fish. For the CAM assay a two-tailed unpaired Student’s t-test was used. The statistical parameters (i.e., exact value of n, p values) have been noted in the figures. Unless otherwise indicated, all data represent the mean ± standard deviation from at least three independent experiments.

### Power calculation

For the *in vivo* experiments the samples size was calculated using a G*Power calculation (Faul et al., 2007). For zebrafish experiments, assuming a difference of 20% in tumorigenic potential and type I error of 5%, 85 samples in each group would ensure >80% power to detect statistical differences between experimental groups using Fisher’s exact test. Furthermore, assuming a 95% engraftment rate, 95 experiments would ensure we had >95% probability of having 85 successful xenotransplantations.

For the CAM assay, assuming an effect size of 1.5 and type I error of 5%, 9 samples in each group would ensure >80% power to detect statistical differences between experimental groups using unpaired t-tests. Furthermore, assuming a 95% engraftment rate, 10 experiments would ensure we had >91% probability of having 9 successful xenotransplantations.

### Data availability

RNA-sequencing data are available at the NCBI Sequence Read Archive (PRJNA623723).

### Code availability

Scripts used to generate the figures for the analysis of the RNA-sequencing are available on request from the corresponding authors.

## Supplemental figure legends

**Figure S1: *GATA3* and *MDM2* are synthetic lethal in ER-positive breast cancer. (A)** Immunoblot showing *GATA3*-mutant, *GATA3*-wild type and MDM2 protein expression levels in MCF-7 cells at 72 h post-siRNA transfection (upper panel). *MDM2* mRNA expression levels (relative to *GAPDH*) in MCF-7 cells at 24, 48 and 72 h post-siRNA transfection (bottom panel). (**B)** Immunoblot showing MDM2 protein expression levels at 72 h post-siRNA transfection (upper panel) and *MDM2* mRNA levels at 48 h post-siRNA transfection (bottom panel) after transfection with different concentrations of siRNA (6.25 nM, 12.5 nM or 25 nM) in MCF-7 cells. (**C)** Proliferation kinetics of MCF-7 cells transfected with *MDM2* siRNA at different concentrations. (**D,E)** *MDM2* and *GATA3* mRNA expression level (relative to *GAPDH*) in (**D)** BT-474 and (**E)** MDA-MB134 cells at 24, 48 and 72 h post-siRNA transfection (left panel). Immunoblot showing *MDM2* and *GATA3* protein levels of expression in (**D)** BT-474 and (**E)** MDA-MB134 cells 72 h post-siRNA transfection (right panel). (**F)** Flow cytometry analysis of Annexin V and propidium iodide co-staining to measure the percentage of apoptotic cells (AnnV+) and live cells (AnnV-/PI-) upon *MDM2* silencing with different concentrations of siRNA in MCF-7 cells. For all the western blots, quantification is relative to the loading control (actin) and normalized to the corresponding siCTR control. Data are mean ± s.d. (**A,B,C,D,E,F)** n≥2 replicates. Statistical significance was determined for (**A,B,C,D,E,F)** by the two-tailed unpaired Student’s t-test.

**Figure S2: *GATA3* status determines response to MDM2 inhibitors *in vitro*. (A)** Logdose response curve of idasanutlin in MCF-7 cells. (**B)** Effect of *GATA3* silencing on proliferation upon treatment with DMSO or idasanutlin 12.5 μM in MDA-MB134 cells. (**C)** logdose response curve of Idasanutlin in MDA-MB134 cells transfected with control siRNA or *GATA3* siRNA. (**D)** Percentage of apoptotic cells upon *GATA3* silencing and Idasanutlin treatment (12.5 μM) in MDA-MB134 cells. (**E)** Immunoblot showing pro- and anti-apoptotic proteins at 12 and 24 h post-treatment with DMSO or Idasanutlin 12.5 μM in MDA-MB134 cells transfected with control siRNAs or *GATA3* siRNAs. For all the western blots, quantification is relative to the loading control (actin) and normalized to the corresponding DMSO control. Data are mean ± s.d. (**A,B,C,D)** n≥3 replicates. Statistical significance was determined for (**B,C,D)** by the two-tailed unpaired Student’s t-test.

**Figure S3: GATA3 expression determines response to MDM2 inhibitor *in vivo***. Representative micrographs of BT-474 tumors extracted four days post-implantation. Tumoural cells (hematoxylin/eosin; upper panel) were immunostained with the apoptotic marker cleaved caspase 3 (lower panel) in the different treatment conditions. Scale bars: 100 μm and 50 μm.

**Figure S4: The synthetic lethality between *GATA3* and *MDM2* is *TP53* dependent. (A)** *MDM2* and *GATA3* mRNA level of expression (relative expression to *GAPDH*) in T-47D at 24, 48 and 72 h post siRNA transfection (left panel). Immunoblot showing *MDM2* and *GATA3* protein level of expression in T-47D cells 72 h post-siRNA transfection (right panel). (**B)** *MDM2* and *TP53* mRNA level of expression (relative to expression of *GAPDH*) in MCF-7 cells 48 h post-siRNA transfection (left panel). Immunoblot showing MDM2 and p53 protein levels of expression in MCF-7 cells 72 h post-siRNA transfection (right panel). (**C)** Immunoblot showing MDM2 and p53 protein levels 24 h post-treatment with DMSO or idasanutlin (12.5 μM) in MCF-7 cells transfected with control siRNAs or *TP53* siRNAs. (**D)** mRNAs levels of *BCL2* and *BAX* in control and *TP53*-silenced MCF-7 cells at 12 and 24 h post-treatment. For all the western blots, quantification is relative to the loading control (actin) and normalized to the corresponding DMSO control. Data are mean ± s.d. (**A,B,D)** n≥2 replicates. Statistical significance was determined for (**A,B,D)** by the two-tailed unpaired Student’s t-test.

**Figure S5: *GATA3* mutations predict response to idasanutlin in patient-derived organoids**. (**A**) Chromatogram obtained from the Sanger sequencing of the *GATA3* gene in the primary (left) and metastatic (right) ER-positive *GATA3*-mutant breast cancer PDOs. (**B**) Representative micrographs of H&E, ER□ and GATA3 immuno-staining on the *GATA3*-mutant (met) PDO. (**C**) Log-dose response curve of idasanutlin in *GATA3* wild-type (IC50=5.4 μM) or *GATA3*-mutant (met) (3.5 μM) PDOs. (**D**) Percentage of viable cells upon treatment with different dosages of idasanutlin in *GATA3* wild-type or *GATA*3-mutant (met) PDOs. Scale bars are 20 and 40 μm for (**B**). Data are mean ± SD, n≥4 replicates from two independent experiments (**C, D**). Statistical significance was determined for (**C,D)** by twotailed unpaired Student’s t-test.

**Figure S6: The synthetic lethality between *GATA3* and *MDM2* acts via the PI3K-Akt-mTOR signaling pathway. (A)** Immunoblot showing MDM2 protein expression in MCF-7 (upper) and MDM2, GATA3, p53, PARP and cleaved PARP protein expression in BT-474 cells (bottom). (**B)** Immunoblot showing markers of mTOR signaling pathway activation at 48 h post siRNA transfection and 24 h post-treatment with DMSO or Idasanutlin in BT-474 cells. Note that this immunoblot was performed using the same membrane as the immunoblot for 24 h in Figure 2G. (**C**) Representative immunohistochemistry of phospho-Akt staining in BT-474 tumors extracted four days post-implantation. Scale bars: 100 μm and 50 μm. (**D,E**) Doughnut charts showing the mutual exclusivity between (**D**) *GATA3* and *PIK3CA* and (**E**) *GATA3* and *PTEN* genetic alterations in ER-positive breast cancer patients. (**F,G**) Normalized enrichment scores of significantly up- and down-regulated pathways identified by gene set enrichment analysis (**F)** in ER-positive breast cancers with low *GATA3* versus high *GATA3* expression and (**G)** in ER-positive breast cancers with *GATA3*-mutant versus *GATA3* wild-type. Statistical significance was determined for (**D,E)** by one-sided Fisher’s Exact test and for (**F,G)** by *fgsea(Sergushichev)*. For all the western blots, quantification is relative to the loading control (actin) and normalized to the corresponding siCTR control or DMSO control.

