## Supplementary figures and images for "GATA3 and MDM2 are synthetic lethal in estrogen receptor-positive breast cancers"

### Supplementary Figures

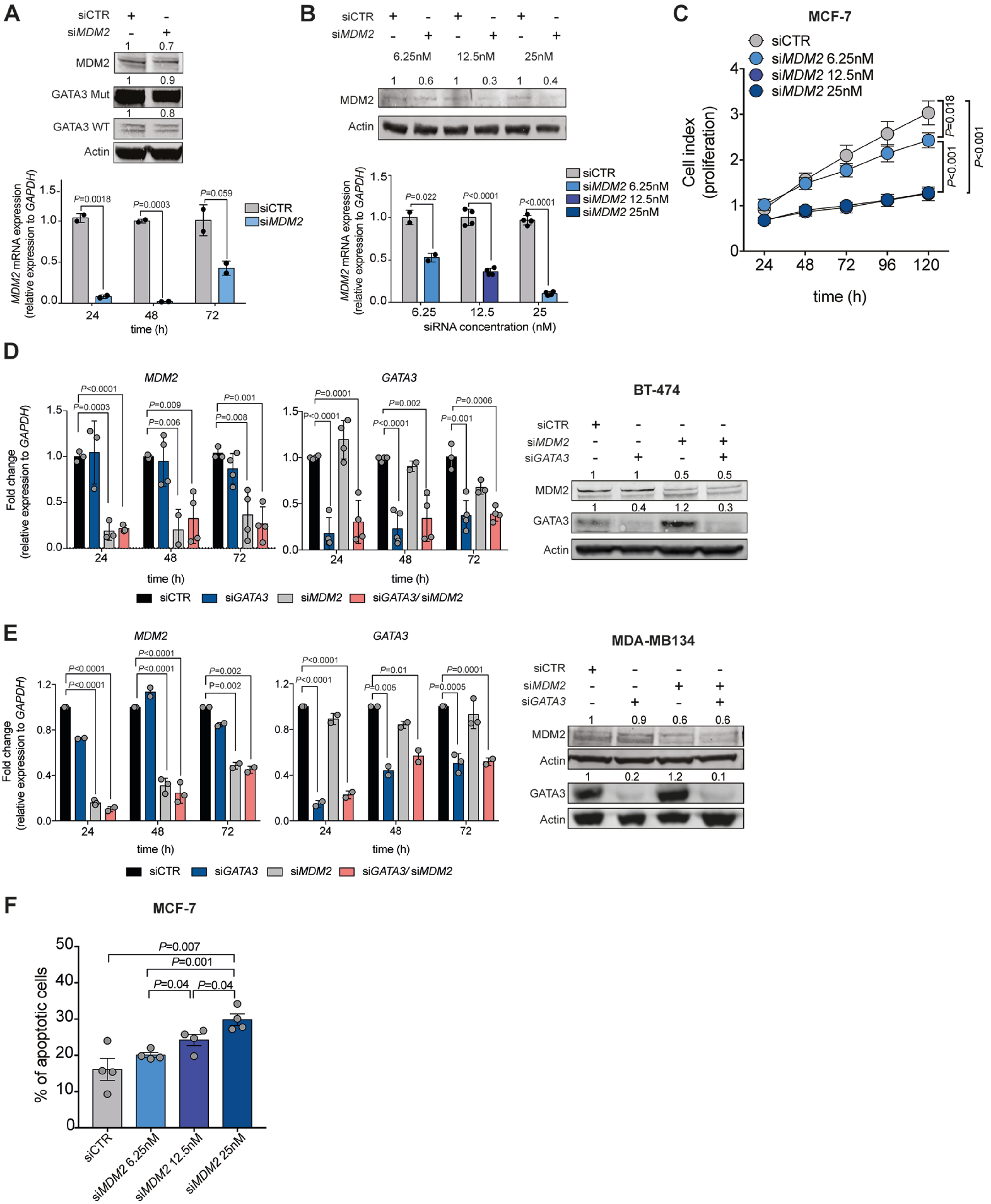

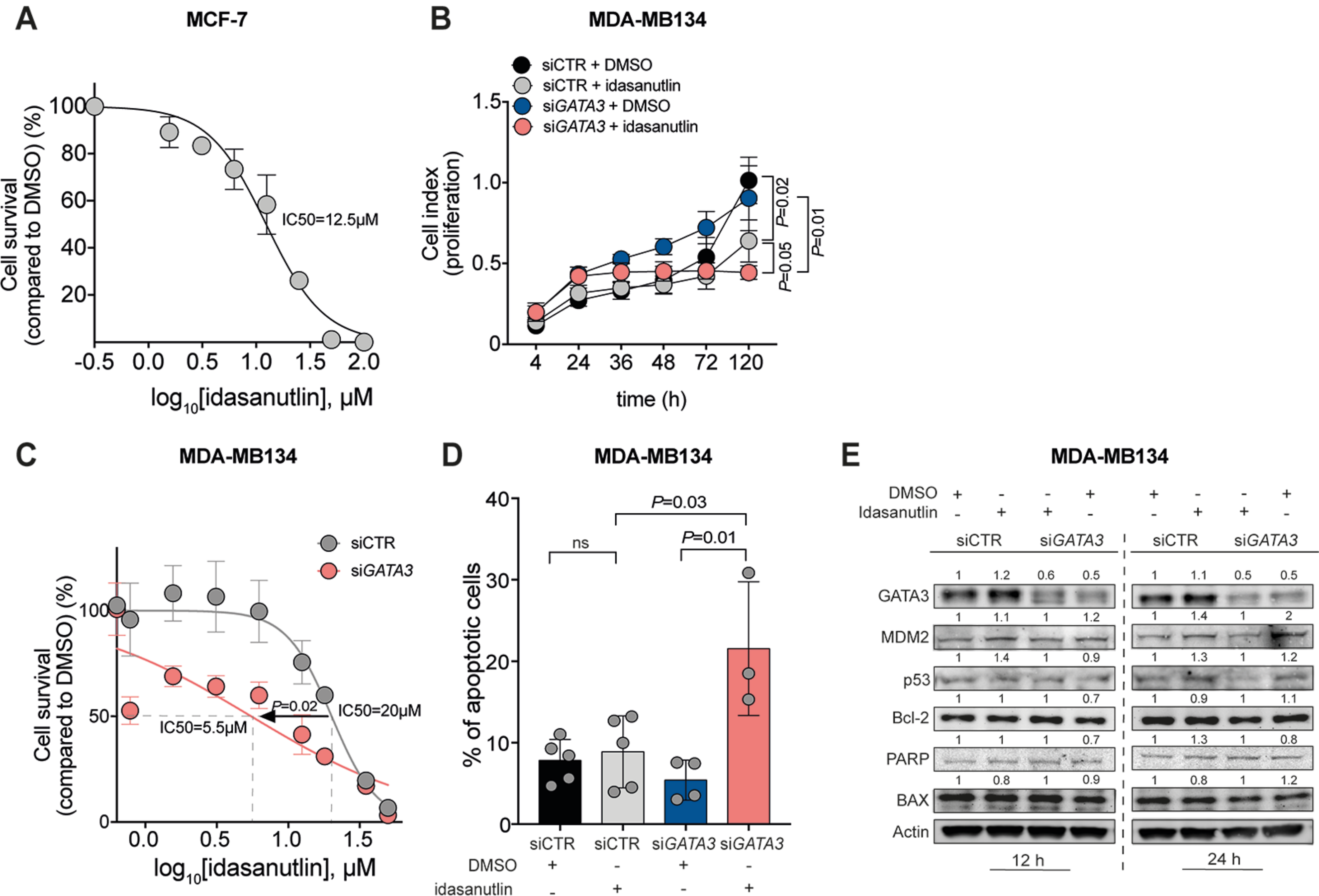

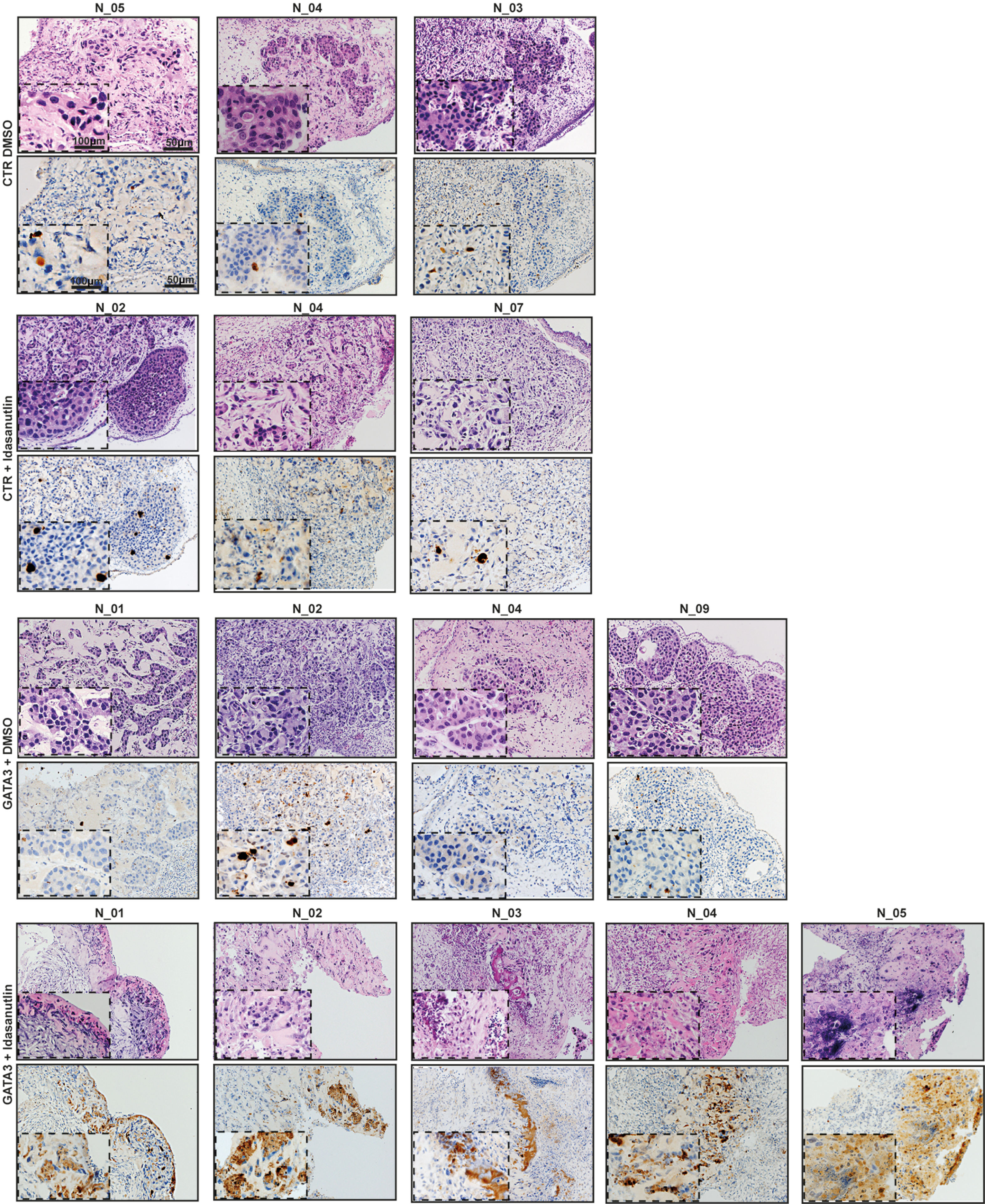

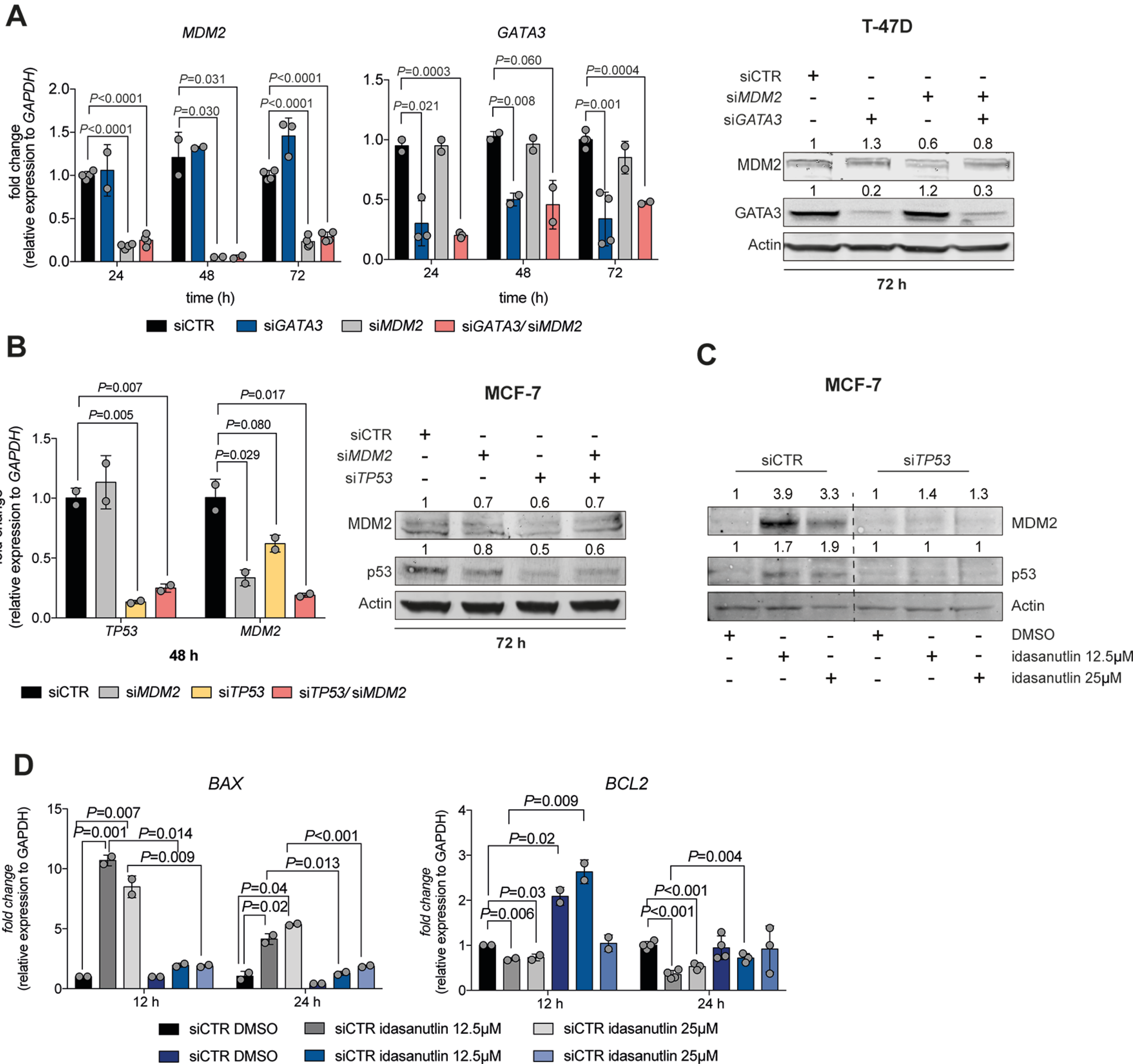

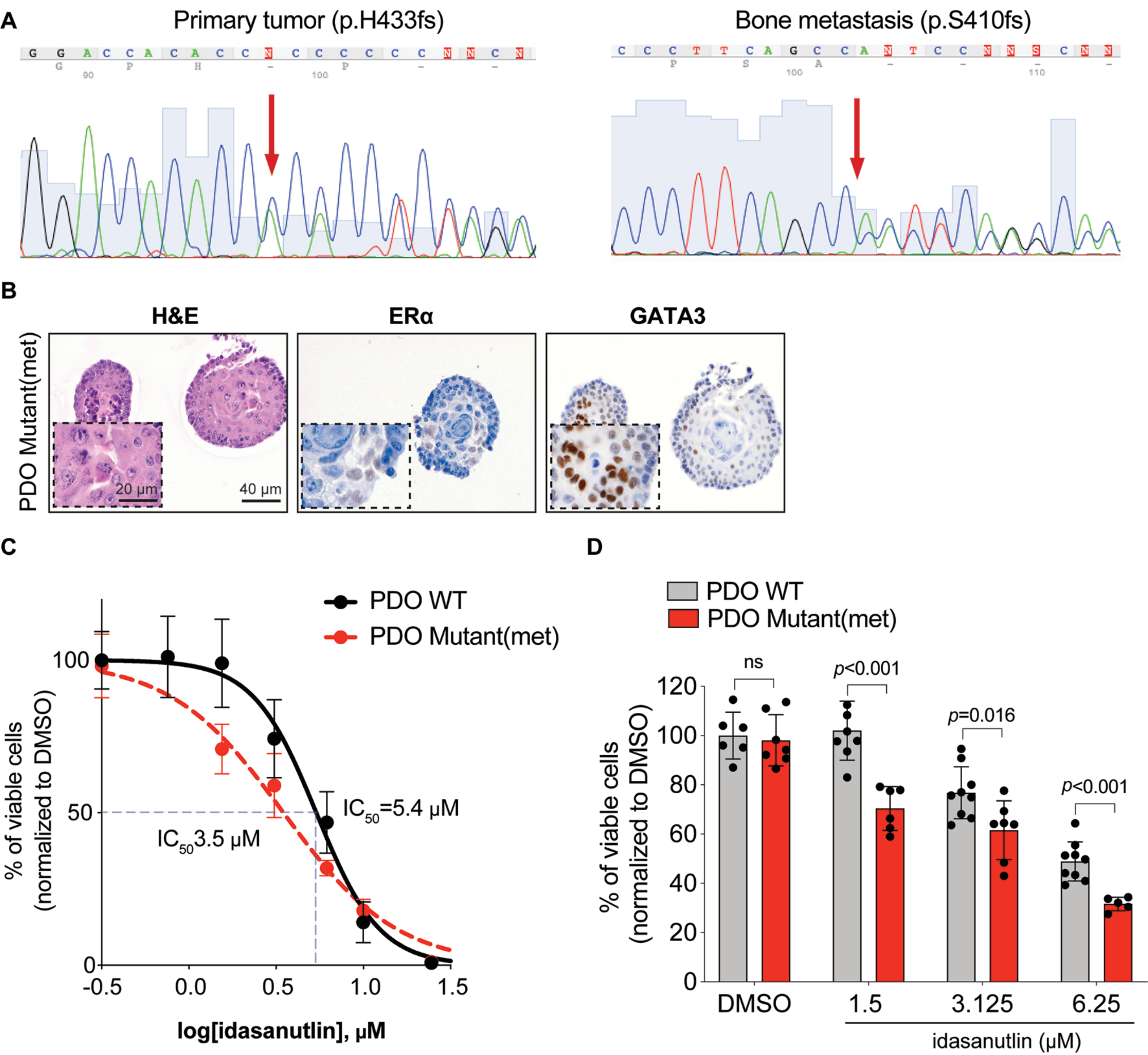

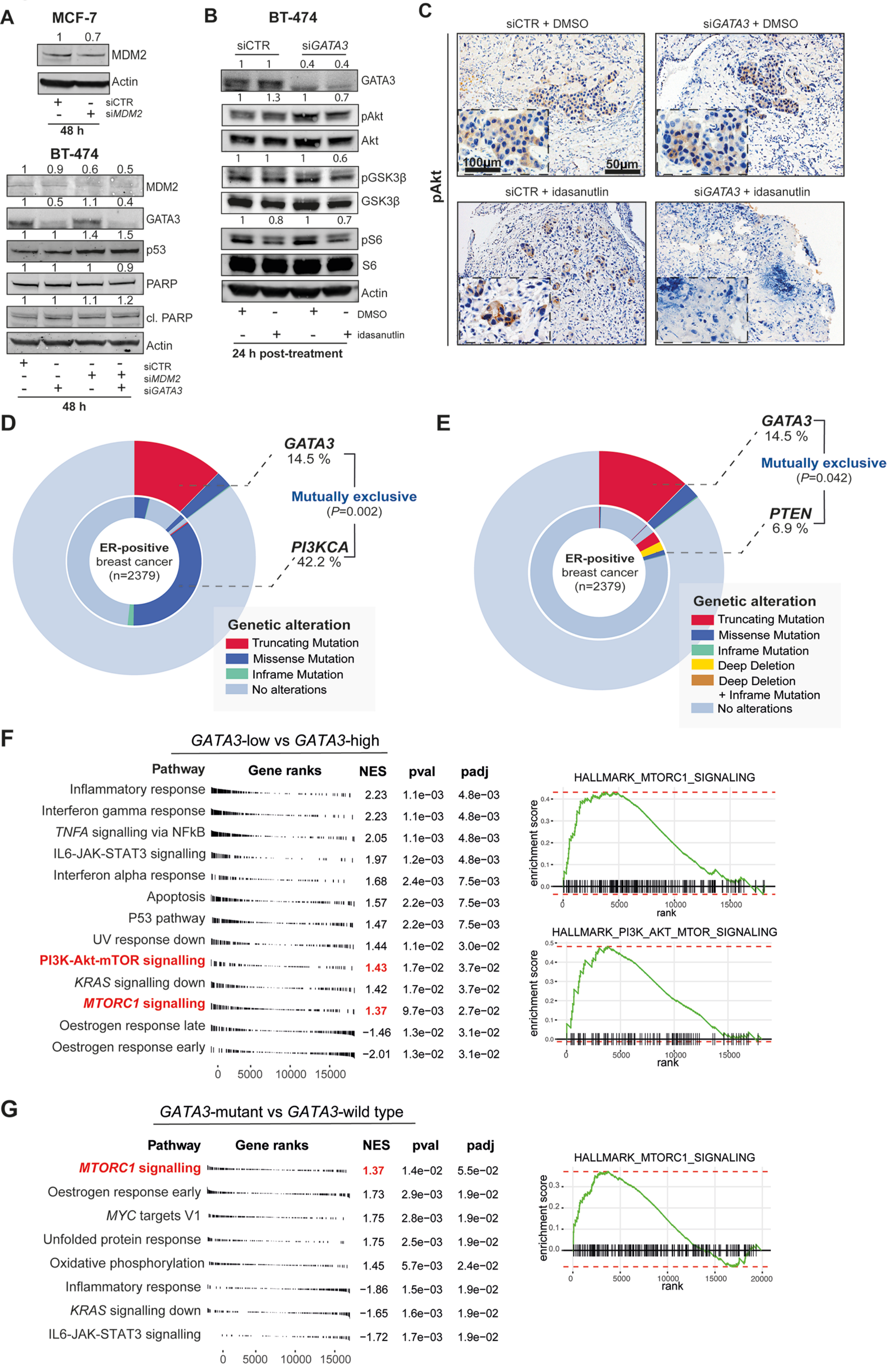
