## Supplementary material for "GATA3 and MDM2 are synthetic lethal in estrogen receptor-positive breast cancers": Key Resources Table

Key Resources Table: List of all the reagents (including antibodies, chemicals, peptides, recombinant proteins, commercial assays, oligonucleotides and qPCR primers)used in this study.

| REAGENT | SOURCE | REFERENCE | NOTES |
|---|---|---|---|
| **Antibodies** | | | |
| **Immunoblot** | | | |
| GATA3 (EPR16651) | abcam | ab199428 | 1 : 1000 |
| MDM2 | MerkMillipore | MABE281 | 1 : 50 |
| p53 (DO-1) | abcam | ab1101 | 1 : 250 |
| BCL-2 | Cell signalling (CST) | 2872S | 1 : 1000 |
| PARP/cl.PARP | Cell signalling (CST) | 9542 | 1 : 1000 |
| BAX (D2E11) | Cell signalling (CST) | 5023 | 1 : 1000 |
| AKT | Cell signalling (CST) | 9272S | 1 : 1000 |
| phospho-AKT (Ser473) (D9E) | Cell signalling (CST) | 4060S | 1 : 1000 |
| S6 ribosomal protein (5G10) | Cell signalling (CST) | 2217S | 1 : 1000 |
| phospho-S6 ribosomal protein (Ser235/236) (D57.2.2E) | Cell signalling (CST) | 4858S | 1 : 2000 |
| GSK-3beta (D5C5Z) | Cell signalling (CST) | 12456S | 1 : 1000 |
| phospho-GSK-3beta (Ser21/9) | Cell signalling (CST) | 9331S | 1 : 1000 |
| Actin | Sigma | A5441 | 1 : 2000 |
| **Immunoistochemistry** | | | |
| GATA3 | abcam | EPR16651 | 1 : 500 |
| Cleaved Caspase 3 (D175) | Cell signalling (CST) | 9661S | 1 : 200 |
| phospho-S6 ribosomal protein (Ser235/236) (D57.2.2E) | Cell signalling (CST) | 4858S | 1 : 400 |
| phospho-AKT (Ser473) (D9E) | Cell signalling (CST) | 4060S | 1 : 100 |
| **Chemicals, Peptides, and Recombinant Proteins** | | | |
| RG7388 (idasanutlin) | Selleckchem | S7205 | N/A |
| DMSO | Sigma-Aldrich | D2650 | N/A |
| Annexin V, FITC conjugate | Invitrogen | V13242 | N/A |
| Propidium iodide (PI) | Invitrogen | V13242 | N/A |
| Annexin V binding buffer 5x | Invitrogen | V13243 | N/A |
| CellTracker™ CM-Dil | Life Technologies | C7000 | N/A |
| cOmplete™, Mini, EDTA-free Protease Inhibitor Cocktail | Roche | 4693159001 | N/A |
| PhosSTOP, Posphatase Inhibitor Cocktail | Roche | 4906845001 | N/A |
| Matrigel® Basement Membrane Matrix | Corning | 354234 | N/A |
| FastStart Universal SYBR Green Master Mix | MerkMillipore | 4913850001 | N/A |
| TRIZOL | Invitrogen | 15596026 | N/A |
| Tricaine methanesulfonate | Sigma-Aldrich | E10521 | N/A |
| MACS Tissue Storage Solution | Miltenyi Biotech | 130-100-008 | N/A |
| DMEM/F-12 | GIBCO | 12634028 | N/A |
| collagenase IV | Worthington | LS004189 | 2.5 mg/mL |
| DNase IV | Sigma | D5025 | 0.1 mg/mL |
| hyaluronidase V | Sigma | H6254 | 20 ug/mL |
| BSA | Sigma | A3059 | 1% |
| LY27632 | Abmole Bioscience | M1817 | 10 µM |
| Accutase | Sigma | A6964 | N/A |
| R-Spondin | Peprotech | 120-38 | 250 ng/ml |
| Neuregulin 1 | Peprotech | 100-03 | 5 nM |
| FGF7 | Peprotech | 100-19 | 5 ng/ml |
| FGF10 | Peprotech | 100-26 | 20 ng/ml |
| EGF | Peprotech | AF-100-15 | 5 ng/ml |
| Noggin | Peprotech | 120-10C | 100 ng/ml |
| A83-01 | Tocris Bioscience | 2939 | 500 nM |
| Y-27632 | Abmole Bioscience | M1817 | 5 mM |
| SB202190 | Sigma-Aldrich | S7076 | 500nM |
| B27 supplement | Life Technologies | 17504044 | 1x |
| N-Acetylcystein | Sigma-Aldrich | A9165 | 1.25 mM |
| Nicotamide | Sigma-Aldrich | N0636 | 5 mM |
| GlutaMax 100x | GIBCO | 35050-068 | 1x |
| Hepes 1M | GIBCO | 15630-056 | 10 mM |
| Penicillin/Streptomycin | GIBCO | 10378016 | 100 mg/ml |
| Primocin | Invivogen | 9155414 | 100 µg/ml |
| Advanced DMEM/F12 | GIBCO | 12634-028 | 1x |
| Trypsin-EDTA | GIBCO | 5200056 | N/A |
| **Commercial Assays** | | | |
| CellTiter-Glo Luminescent Cell Viability Assay | Promega | G7570 | N/A |
| CellTiter-Glo 3D Luminescent Cell Viability Assay | Promega | G9682 | N/A |
| SuperScript™ VILO™ cDNA Synthesis Kit | Invitrogen | 11754050 | N/A |
| ON-TARGET plus siRNA transfection reagent | Dharmacon | T-2001-03 | N/A |
| **Oligonucleotides** | | | |
| **siRNA** | | | |
| SMARTpool siRNAs against human *GATA3* | Dharmacon | L-003781-00-0005 | N/A |
| SMARTpool siRNAs against human *MDM2* | Dharmacon | L-003279-00-0005 | N/A |
| SMARTpool siRNAs against human *TP53* | Dharmacon | L-009625-00-0005 | N/A |
| ON-TARGET plus SMARTpool non-targeting control | Dharmacon | D-001810-10-05 | N/A |
| **qRT-PCR primers** | | | |
| *GATA3*_Forward (TCGCAGAATTGCAGAGTCGT) | | | |
| *GATA3*_Reverse (GAGTTTCCGTAGTAGGGCGG) | | | |
| *MDM2*_Forward (GGCGAGCTTGGCTGCTTC) | | | |
| *MDM2*_Reverse (TGAGTCCGATGATTCCTGCTG) | | | |
| *TP53*_Forward (TGCTCAAGACTGGCGCTAAA) | | | |
| *TP53*_Reverse (TTTCAGGAAGTAGTTTCCATAGGT) | | | |
| *BCL2*_Forward (TCTTTGAGTTCGGTGGGGTC) | | | |
| *BCL2*_Reverse (GACTTCACTTGTGGCCCAGAT) | | | |
| *BAX*_Forward (GCCCTTTTCTACTTTGCCAGC) | | | |
| *BAX*_Reverse (AGACAGGGACATCAGTCGC) | | | |
| *PUMA*_Forward (CTGCCAGATTTGTGGTCCTC) | | | |
| *PUMA*_Reverse (CCTTCCGATGCTGAGTCCAT) | | | |
| GAPDH83U (AGGTGAAGGTCGGAGTCAACG) | | | |
| GAPDH28L (TGGAAGATGGTGATGGGATTT) | | | |
